# Developmental priming of cancer susceptibility

**DOI:** 10.1101/2023.09.12.557446

**Authors:** Ilaria Panzeri, Luca Fagnocchi, Stefanos Apostle, Megan Tompkins, Emily Wolfrum, Zachary Madaj, Galen Hostetter, Yanqing Liu, Kristen Schaefer, Yang Chih-Hsiang, Alexis Bergsma, Anne Drougard, Erez Dror, PERMUTE, Darrell Chandler, Daniel Schramek, Timothy J. Triche, J. Andrew Pospisilik

## Abstract

DNA mutations are necessary drivers of cancer, yet only a small subset of mutated cells go on to cause the disease. To date, the mechanisms that determine which rare subset of cells transform and initiate tumorigenesis remain unclear. Here, we take advantage of a unique model of intrinsic developmental heterogeneity (*Trim28^+/D9^*) and demonstrate that stochastic early life epigenetic variation can trigger distinct cancer-susceptibility ‘states’ in adulthood. We show that these developmentally primed states are characterized by differential methylation patterns at typically silenced heterochromatin, and that these epigenetic signatures are detectable as early as 10 days of age. The differentially methylated loci are enriched for genes with known oncogenic potential. These same genes are frequently mutated in human cancers, and their dysregulation correlates with poor prognosis. These results provide proof-of-concept that intrinsic developmental heterogeneity can prime individual, life-long cancer risk.

## INTRODUCTION

Cancer is triggered by oncogenic DNA mutations^1–3^. However, these mutations are also found at relatively high rates in otherwise ‘normal’ tissues, and not every mutation is oncogenic across all tissues^4–13^. In other words, the oncogenic potential of DNA mutations are cell-, tissue-, and temporal-specific^14,15^. The molecular basis of this context specificity comprises one of the biggest unanswered questions in cancer biology.

Pioneering studies over the last decades have implicated epigenetic regulation as a key mediator of context specificity. Notable examples include demonstrations that cell-type and differentiation-stage specific differences in epigenetic control determine when and where transformation occurs^14,16–23^. What is typically overlooked in human genetics and epidemiology is that, in addition to the epigenetic programs that emerge and drive cell differentiation, another layer of *intrinsic* epigenetic variation arises during development that is at least partially *stochastic* in nature^24^. Indeed, these epigenetic changes occur at rates several orders of magnitude higher than mutations^25^, and an unequal distribution of epigenetic marks can drive phenotypic discordance for instance between MZ twins or isogenic mice^25–27^. This ‘**intrinsic developmental heterogeneity**’ is distinct from the epigenetic changes triggered by *external* environmental exposures and from the large literature of *in utero* and early-life environmental insults that can increase cancer risk (e.g., estrogens, pesticides, alcohol, and over-or under-nutrition^28–33^). While an impressive theoretic framework has been developed for how intrinsic developmental heterogeneity impacts cancer^34^, to our knowledge, the notion has never been demonstrated experimentally. This has been due in part to a lack of proper models.

Because tumor initiation involves some degree of randomness, testing the relationship between developmental heterogeneity and cancer susceptibility requires measurement of the *distribution* of observed outcomes comparing distinct intrinsic epigenetic states; essentially, it requires an isogenic model, raised in tightly controlled environments, but bearing more than one reproducible intrinsic epigenetic states^35,36^. TRIM28 is an epigenetic regulator that plays an important role in heterochromatic gene silencing^37–41^.TRIM28 loss-of-function models have implicated the protein in cancer in complex and tissue-specific manner^42^. TRIM28, however, is also a master regulator of organism-level developmental heterogeneity^43^. Our prior work showed that genetically and environmentally identical *Trim28^+/D9^* haplo-insufficient mice emerge into adulthood as two distinct populations (or developmental morphs) characterized by differences in body mass composition^44^. The *Trim28^+/D9^* mouse thus meets the requirements for testing intrinsic developmental heterogeneity effects; it provides a model sensitized to detect the long-term phenotypic consequences of two distinct developmental programming states. Here, we leverage this unique model and ask if isogenic populations with reproducibly distinct intrinsic epigenetic states might exhibit differential cancer susceptibility. We show that the two *Trim28^+/D9^* developmental morphs develop distinct types, timing and severity of cancer. We identify a signature of DNA hypo-methylated genes, installed well before weaning, that stratify mice for cancer risk and outcome. These same genes are frequently mutated in human cancers and their dysregulation correlates with poor prognosis, suggesting that if conserved, this novel mode of action has the potential to impact a broad portion of the population.

## RESULTS

### *Trim28^+/D9^* mice exhibit a novel multi-cancer syndrome

To test if developmental heterogeneity influences cancer, we crossed B6J.*Tp53^+/R270H^*mice with FVB.*Trim28^+/D9^* animals (**Fig.1A**). The *Tp53^+/R270H^*mouse is a multi-cancer syndrome (MCS) model^45^, while the *Trim28^+/D9^* mouse is sensitized to exhibit reproducible and exaggerated developmental heterogeneity^43,44,46,47^. Both lines were highly backcrossed and cohorts thus yielded isogenic offspring with one of four genotypes: wild-type (*WT*), *Trim28^+/D9^* single heterozygotes (*Trim28*), *Tp53^R270H/+^* single heterozygotes (*Tp53*), and *Tp53^R270H/+^*;*Trim28^+/D9^* compound heterozygotes (*Tp53/Trim28*). Parental ID, litter-size, and housing density were all carefully recorded to maximize our ability to associate cancer outcomes with developmental heterogeneity while reducing confounders. The experiment tracked animals from birth to endpoint (70 weeks of age), monitoring every individual for signs of cancer 2-3 times per week, with periodic measures of morphological, growth, and metabolic characteristics (**Fig.1A**). Early-life ear biopsies were obtained at 10 days of age for epigenomic profiling (**Fig.1A**). At sacrifice, all animals underwent a systematic 21-organ dissection protocol in which tissues were isolated, processed for histology, and scored by a pathologist. Cancer events were divided into *aggressive* (i.e., animals requiring euthanasia prior to 70 weeks) and *endpoint* events (i.e., animals reaching the 70-weeks endpoint without evidence of sickness). The final dataset comprised 137 animals with 79 malignant and 34 benign primary tumors.

**Figure 1.**
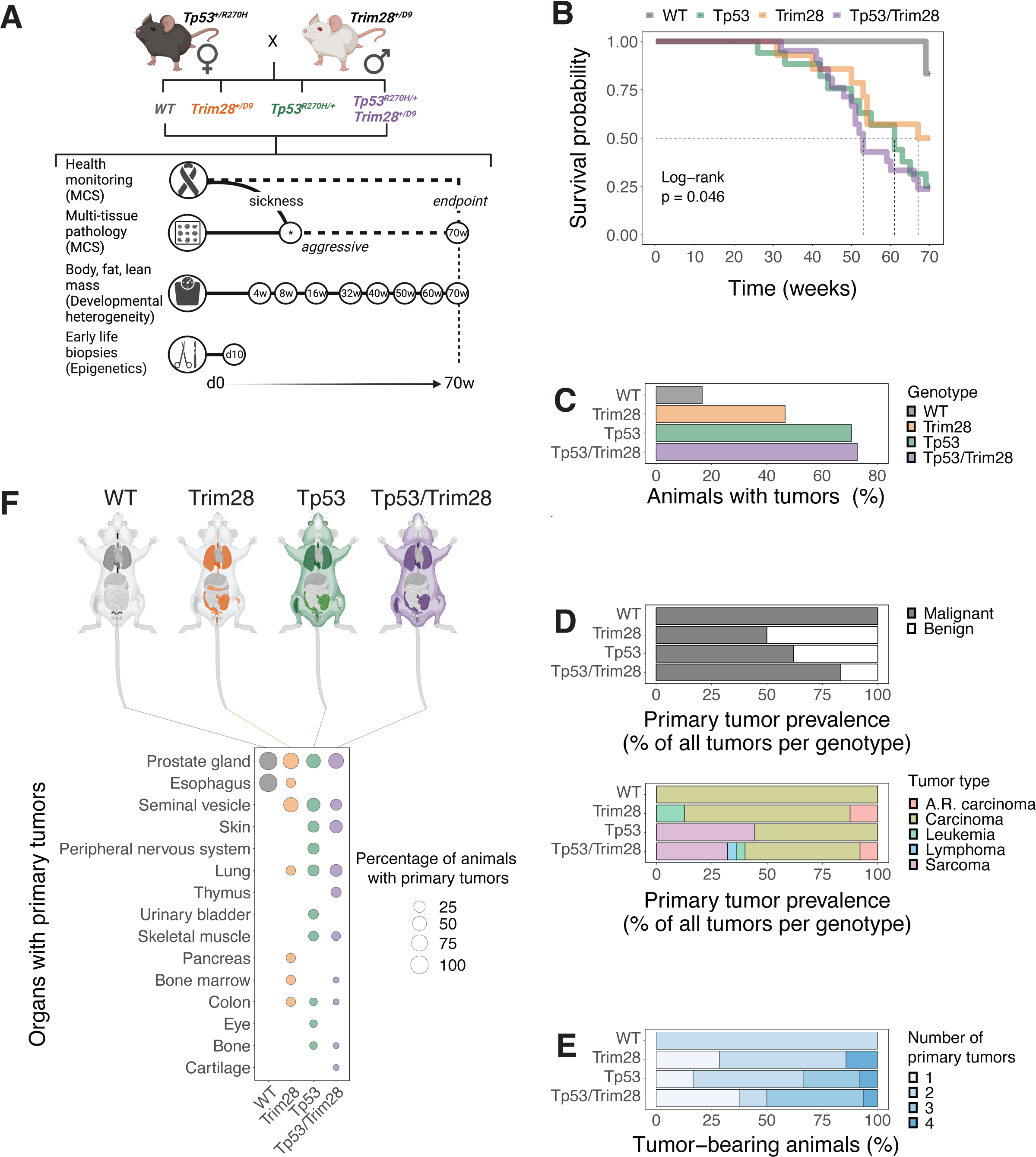
*Trim28^+/D^*^9^ mice exhibit a novel multi-cancer syndrome. **A)** Schematic of the experimental plan. We mated the *Tp53^+/R270H^* multi-cancer syndrome model (MCS) with the *Trim28^+/D9^* developmental heterogeneity model. All the resulting genotypes were screened for health issues and tumor development. Tissues and masses were harvested either before the endpoint (*aggressive*) or at the *endpoint* of the study (70 weeks). Histopathological evaluation was performed on the harvested tissues and masses to determine the presence of tumors. Body, fat, and lean mass composition data were collected at multiple timepoints. Early life biopsies were collected at day 10 of age (pre-weaning). Created with BioRender.com. **B)** Kaplan-Meier survival probability for each genotype. Log-rank test, p=0.046. N=60 (6 WT, 17 *Tp53^R270H/+^*, 15 *Trim28^+/D9^*, 22 *Tp53^R270H/+^;Trim28^+/D9^*). **C)** Percentage of animals found with *aggressive* tumors for each genotype (relative to the total number of animals screened for each genotype). N=60 (6 WT, 15 *Trim28^+/D9^*, 17 *Tp53^R270H/+^*, 22 *Tp53^R270H/+^;Trim28^+/D9^*). **D)** <u>Top panel</u>: prevalence of *malignant* (black bars) or *benign* (white bars) *aggressive* tumors of for each genotype, expressed as percentage relative to the total number of tumors found in each genotype. N=76 (total tumors: 1 in WT, 16 in *Trim28^+/D9^*, 29 in *Tp53^R270H/+^*, 30 in *Tp53^R270H/+^;Trim28^+/D9^*). <u>Bottom panel</u>: prevalence of distinct *malignant aggressive* tumor types for each genotype, expressed as percentage relative to the total number of *malignant aggressive* tumors found in each genotype. N=52 (total malignant aggressive tumors: 1 in WT, 8 in *Trim28^+/D9^*, 18 in *Tp53^R270H/+^*, 25 in *Tp53^R270H/+^;Trim28^+/D9^*). **E)** Fraction of animals harboring 1 or multiple *aggressive malignant* tumors prior to endpoint in the different genotypes. N=36 animals sacrificed prior to endpoint (1 WT, 7 *Trim28^+/D9^*, 12 *Tp53^R270H/+^*, 16 *Tp53^R270H/+^;Trim28^+/D9^*). **F)** Tissues targeted by *malignant aggressive* tumors in the different genotypes. <u>Top panel</u>: mouse anatomy plots, with non-targeted in light-grey and targeted tissues colored according to the affected genotype: WT in black, *Trim28^+/D9^* in orange, *Tp53^R270H/+^* in green, *Tp53^R270H/+^*;*Trim28^+/D9^* in purple. <u>Bottom panel</u>: percentage of animals with specific organs targeted by *malignant aggressive* tumors in the different genotypes. N=36 animals sacrificed prior to endpoint (1 WT, 7 *Trim28^+/D9^*, 12 *Tp5^R270H/+^*, 16 *Tp53^R270H/+^;Trim28^+/D9^*). Created with BioRender.com

As expected^45,48,49^, *Tp53^R270H/+^* siblings exhibited a high penetrance multi-cancer syndrome (MCS), with 76% of *Tp53^R270H/+^* mice succumbing to aggressive tumors before the 70-week endpoint (**Fig.1B-C**). We found 24 total primary malignant tumors in *Tp53^R270H/+^* animals, primarily consisting of carcinomas and sarcomas. Eighteen of these developed before the 70-week study endpoint (**Fig.1C**) and were widely distributed throughout the body (**Fig.1D-F** and **S1E**). We were surprised to find that *Trim28^+/D9^* heterozygotes also showed a substantially reduced survival probability, similar to that of *Tp53^R270H/+^*animals (**Fig.1B**; mean survival probability of 59.9 and 56.5 weeks, respectively). Health-monitoring and the 21-organ histopathology analysis revealed that reduced *Trim28^+/D9^* survival was due to its own MCS. *Trim28^+/D9^* animals showed similar time-to-detection and tumor burden to *Tp53^R270H/+^* animals (**Fig.1B, 1D**, top, and **-1E**). That said, *Trim28^+/D9^*-triggered MCS showed several differences relative to *Tp53^R270H/+^*-triggered MCS. First, *Trim28^+/D9^* animals specifically developed rare germ-cell tumors (**Fig.S1B-C**). Second, *Trim28^+/D9^* animals only showed one case of sarcoma, whereas sarcomas were common in *Tp53^R270H/+^* animals (**Fig.1D**, bottom, and **S1C**). Overall, carcinomas dominated the landscape of malignant primary tumors across genotypes, with representative histological examples shown in **Appendix 1**. *Tp53^R270H/+^;Trim28^+/D9^* compound heterozygotes showed the lowest survival probability of all genotypes (**Fig.1B**), with target tissue and tumor type distributions consistent with the presence of both alleles (**Fig.1F** and **1D**). Stratifying the data into *aggressive* and *endpoint* samples indicated that the early and late pathologies within each genotype were largely constant (**Fig.1A-E** and **S1A-E**). As expected, age-associated carcinomas were over-represented in WT animals (**Fig.S1C** and **S1E**, right). The few sarcomas that were observed in *Trim28^+/D9^* animals were found at endpoint (**Fig.S1C**). Collectively, these data demonstrate that *Trim28^+/D9^* triggers a novel MCS, similar in timing and penetrance to *Tp53^R270H/+^*.

### TRIM28-dependent developmental heterogeneity primes cancer outcomes

Consistent with the literature^43,44,46,47^, *Trim28^+/D9^* animals showed marked variation in body mass at 16 weeks of age (**Fig.2A**) and separated statistically (MClust) into two distinct developmental ‘morphs’ (reproducible phenotypic forms), referred to here as *Trim28^+/D9^*-***heavy*** and *Trim28^+/D9^****-light*** (**Fig.2B-C, S2A-**). *Tp53^R270H/+^*;*Trim28^+/D9^* compound heterozygotes also showed high variation in body mass, indicating that the developmental heterogeneity effect of *Trim28^+/D9^* is maintained in the presence *Tp53^R270H/+^*. In compound heterozygotes, however, bimodality could not be statistically resolved (**Fig.2A-C**, and **Fig.S2A-B**). Neither *WT* nor *Tp53^R270H/+^* heterozygotes showed significant variation phenotypes (**Fig.2A-B**), nor of a bimodal distribution (**Fig.2A-C**, and **Fig.S2A-B**). We validated the unbiased developmental morph calling using Rmixmod and found 100% congruence with MClust (cluster detection and morph assignment) (**Fig.S2C**). The phenotypic distinctions between morphs are transient over the long-term (**Fig.S2A**). They were also consistent with our previous findings^44^. This developmental bifurcation in *Trim28^+/D9^* animals is a critical feature of the model, because it enables comparison of cancer outcomes between groups of isogenic animals with distinct yet *reproducible* developmental trajectories (**Fig.2B** and **Fig.S2A**).

**Figure 2.**
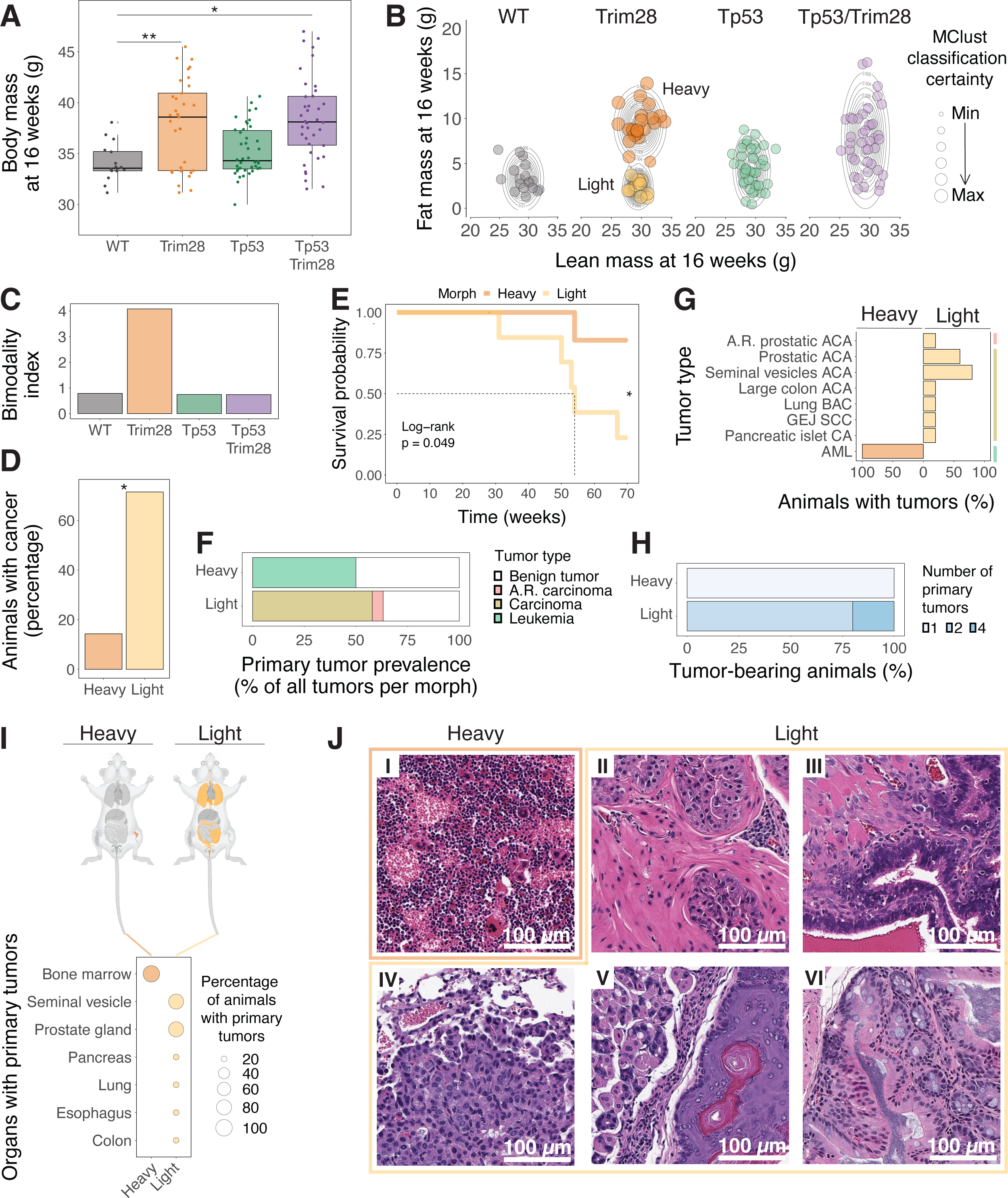
TRIM28-dependent developmental heterogeneity primes cancer outcomes. **A)** Distribution of body mass at 16 weeks of age for each genotype. Each dot represents one animal. Levene’s test, Benjamini–Hochberg adjusted p-value for multiple comparisons, threshold of significance for padj<0.05: * p<0.05; ** p<0.01. WT vs *Trim28^+/D9^*, padj=0.0016; WT vs *Tp53^R270H/+^*, padj=0.2174; WT vs *Tp53^R270H/+^;Trim28^+/D9^*, padj=0.0152; *Trim28^+/D9^* vs *Tp53^R270H/+^*, padj=0.0016; *Trim28^+/D9^* vs *Tp53^R270H/+^;Trim28^+/D9^*, padj=0.4220; *Tp53^R270H/+^;Trim28^+/D9^* vs *Tp53^R270H/+^*, padj=0.0178. N=129 (16 WT, 33 *Trim28^+/D9^*, 42 *Tp53^R270H/+^*, 38 *Tp53^R270H/+^;Trim28^+/D9^*). **B)** Fat and lean mass data at 16 weeks for the different genotypes. Each dot represents one animal. Overlaid density estimation of the data by MClust for each genotype. The dot size represents the classification certainty (probability) as calculated by MClust for each animal. N=137 (18 WT, 34 *Trim28^+/D9^*, 44 *Tp53^R270H/+^*, 41 *Tp53^R270H/+^;Trim28^+/D9^*). **C)** Bimodality index determined for each genotype as the ratio between the MClust-determined Bayesian Information Criterion (BIC) for 2 clusters over 1 cluster from fat and lean mass data at 16 weeks (same data as in Fig.2A). **D)** Proportion of *Trim28^+/D9^*-*heavy* and -*light* animals affected by *malignant aggressive* tumors (expressed as percentage of animals developing tumors relative to the total animals in each population). Two-sample test for equality of proportions without continuity correction, significance for p<0.05: -*heavy* vs -*light*, p=0.0308, χ^2^=4.6667: * p<0.05. N=14 (7 -*heavy*, 7 -*light*). **E)** Kaplan-Meier survival probability of *Trim28^+/D9^*-heavy and -light animals. Log-rank test, p=0.049: * p<0.05. N=14 (7 -*heavy*, 7 -*light*). **F)** Prevalence of distinct *aggressive* tumor types in *Trim28^+/D9^*-*heavy* and -*light* animals (expressed as percentage relative to the total number of *aggressive* tumors found in each population). N=21 (total *aggressive* tumors, including *malignant* and *benign*, 2 in -*heavy* and 19 in -*light*). **G)** Distribution of *malignant aggressive* tumor types in *Trim28^+/D9^*-*heavy* and -*light* animals, expressed as percentage of animals with a particular tumor type relative to the total number of animals with *malignant aggressive* tumors in each population. The colored bars on the right identify the main tumor type, as in Fig.2F: age-related carcinoma in red; carcinoma in gold; leukemia in green. N=6 (1 -*heavy*, 5 -*light*). **H)** Fraction of *Trim28^+/D9^*-*heavy and* -*light* animals that died before the endpoint of the study harboring 1 or multiple *malignant aggressive* tumors. N=6 (1 -*heavy*, 5 -*light*). **I)** Tissues targeted by *malignant aggressive* tumors in the different genotypes. <u>Top panel</u>: mouse anatomy plots, with non-targeted in pale grey and targeted tissues colored according to the affected morph: dark orange for -*heavy*, pale orange for -*light*. <u>Bottom panel</u>: percentage of animals with specific organs targeted by *malignant aggressive* tumors in the different morphs. N=6 (1 -*heavy*, 5 -*light*). **J)** Key histological examples of *aggressive malignant* tumors and targeted tissues for -*heavy* and -*light* animals. I-AML targeting the bone marrow in a *Trim28^+/D9^*-*heavy* animal (same as Appendix 1B-IX). II-Prostatic ACA in a *Trim28^+/D9^*-*light* animal. III-Seminal vesicles ACA *Trim28^+/D9^*-*light* animal. IV-Bronchoalveolar CA in a *Trim28^+/D9^*-*light* animal. V-Gastroesophageal junction SCC in a *Trim28^+/D9^*-*light* animal. VI-Colon ACA in a *Trim28^+/D9^*-*light* animal.

Because *Trim28^+/D9^* animals exhibited both a reproducible developmental bifurcation and a novel MCS, we focused on the *Trim28^+/D9^* genotype to test for associations between intrinsic developmental heterogeneity and cancer outcomes. Using the unbiased MClust assignment of *Trim28^+/D9^* animals to their *-light* or *-heavy* morph, we compared survival, tumor incidence and associated outcomes between the two groups (**Fig. 2D-J**). Whereas 86% of *Trim28^+/D9^-heavy* animals reached the study endpoint illness-free, the majority of *Trim28^+/D9^-light* animals showed aggressive MCS (**Fig. 2D-J**). Mean survival times were significantly different between *Trim28^+/D9^-light* and *-heavy* morphs (55.4 and 67.3 weeks, respectively) (**Fig.2E**). The single *Trim28^+/D9^-heavy* animal that required pre-endpoint euthanasia had a bone-marrow-derived acute myeloid leukemia (AML; **Fig. 2E-I**, and **2J** panel I). In contrast, *Trim28^+/D9^-light* animals exhibited up to four different primary tumors per animal (**Fig. 2H**), mainly consisting of carcinomas, age-related carcinomas, and benign tumors (**Fig 2F**). Tumors in *Trim28^+/D9^-light* animals were found in seminal vesicles, prostate, pancreas, lungs, esophagus, and colon (**Fig. 2G, 2I** and **2J** panels II-VI). Histopathology of the *Trim28^+/D9^-heavy* AML and representative *Trim28^+/D9^-light* carcinomas are shown in **Fig. 2J** and **Appendix 1B**. Consistent with the differential timing of cancer onset between -*light* and *-heavy* animals, the endpoint analysis was dominated by -*heavy* tumors (**Fig.S2D-H**). Thus, *Trim28^+/D9^-light* developmental morphs exhibit an accelerated MCS. These data provide genetic evidence that TRIM28-dependent epigenetic variation in development controls later-life cancer outcomes.

### *Trim28^+/D9^*-dependent cancer susceptibility states are distinguished by distinct early-life epigenomes

We reasoned that if developmentally programmed epigenetic differences impact cancer susceptibility and outcomes late in life, then these differences should be detectable early on. We therefore performed DNA-methylation profiling on biopsies taken from all genotypes and animals at 10 days of age (i.e., before weaning). We used ear clips as they are minimally invasive for young mice and are similar to tissues used to identify early-life epigenetic signatures in humans^50–55^. We used Illumina Infinium Mouse Methylation BeadChips to quantify DNA methylation state at ∼285,000 CpG sites that included CpGs in all annotated genes, functional RNAs, and cis-regulatory regions of the mouse genome. Global DNA methylation profiles were highly correlated across genotypes, indicating a robust technical approach and high sample quality (**Fig. S3A**, row 1-4). *Trim28^+/D9^* and *Tp53^R270H/+^;Trim28^+/D9^* compound heterozygotes had ∼3 times as many differential methylated CpG loci (DML) than *Tp53^R270H/+^* alone (**Fig. S3B**). DMLs in *Trim28^+/D9^* and *Tp53^R270H/+^;Trim28^+/D9^* animals overlapped strongly (**Fig. S3A-3D**), indicating that *Trim28^+/D9^* substantially and reproducibly changes the early life methylome even in the presence of the *Tp53^R270H/+^*mutation.

Relative to WT, *Trim28^+/D9^* biopsies were skewed towards reduced methylation (**Fig. S3E**). This is consistent with TRIM28’s known role in gene silencing ^37,38,40,41,56^. Interestingly, *Tp53^R270H/+^* animals also showed early-life epigenetic changes, and these showed some similarity to *Trim28^+/D9^*-induced changes (**Fig. S3F**). These data are consistent with reports that TP53 regulates DNA methylation^57–59^. *Trim28^+/D9^* hypomethylated DMLs were enriched almost exclusively in regions known to be targeted by TRIM28, with probe set enrichment analysis (PSEA) revealing annotations for *heterochromatin*, *monoallelic methylation*, *Polycomb-silencing*, *CTCF*, *TRIM28-binding*, and *H3K9me3* (**Fig. S3G**). These data show that full TRIM28 dosage is required to maintain early life DNA methylation fidelity at heterochromatic regions.

Importantly, substantial DNA methylation differences were also detected between isogenic *Trim28^+/D9^* animals that would go on to become *-light* versus *-heavy*, even though there are no detectable phenotypic differences at 10 days of age (**Fig. 3A**, **S3A** row 5-6, and **S3H-I**). We found a total of 1133 DMLs between the two *Trim28^+/D9^* morphs, and a clear skew towards hypomethylation in the -*light*, *cancer-prone* morph (**Fig. 3A**, **S3I**). Similar to the genotype as a whole, PSEA revealed differential methylation predominantly at regions of monoallelic methylation and imprinting, including the *Kcnq1-Kcnq1ot1* cluster, *H19*, and *Peg3* (**Fig. 3B** top panel, green, and bottom panel, pale orange), and regions annotated as H3K9me3- and H3K27me3-decorated (**Fig. 3B** top panel, light blue). A search for overlap with transcription factor binding (**Fig. 3B** top panel, purple) revealed strong and specific enrichment for chromatin binders involved in DNA-methylation (MBD1, MECP2, C17orf96, DPPA2 and TRIM28 itself) and Polycomb silencing machinery (SUZ12, EZH2, C17orf96, RNF2, AEBP2, PCGF2, CBX7, BMI1, and JARID2).

**Figure 3.**
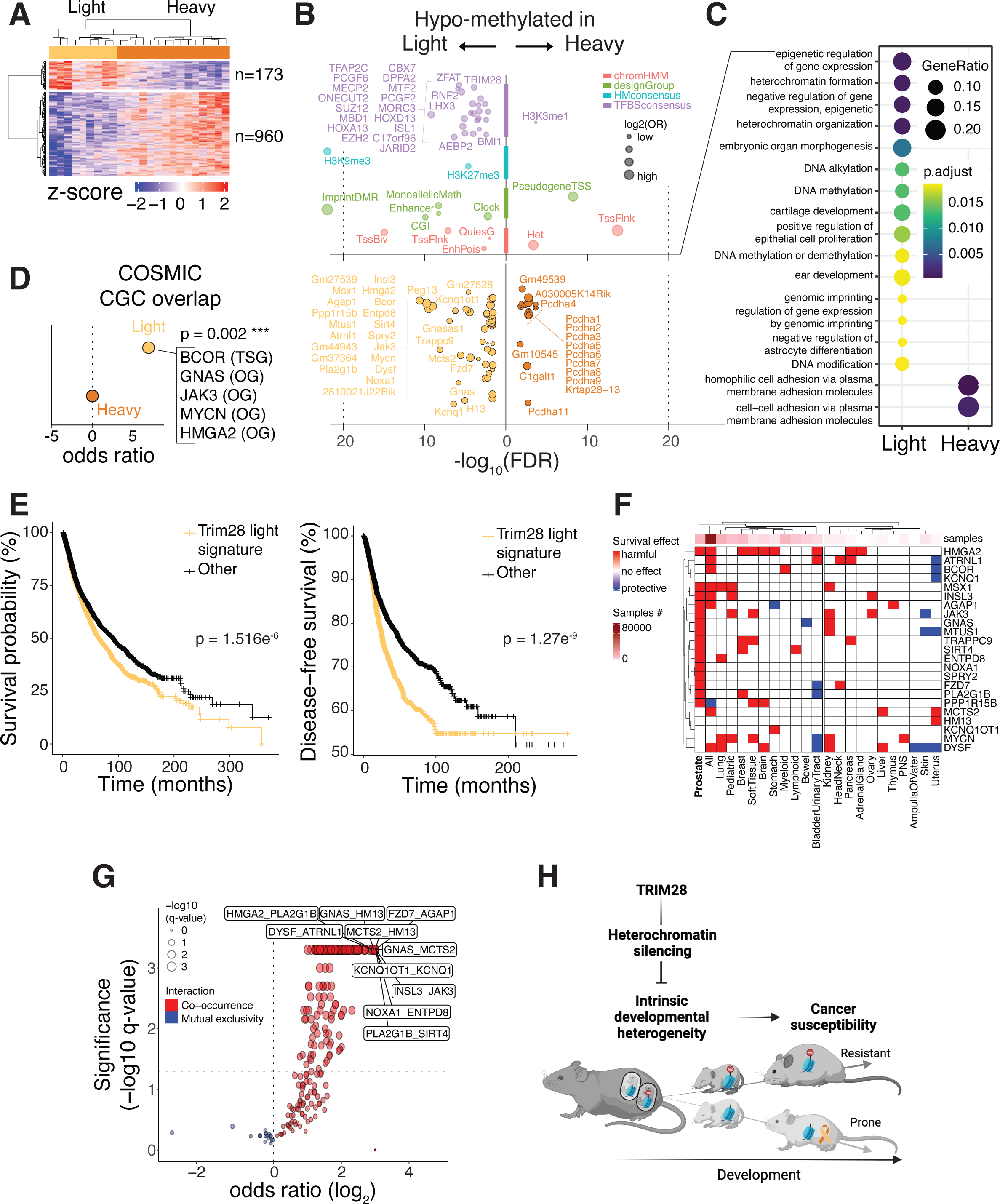
*Trim28^+/D9^*-dependent cancer susceptibility states are distinguished by distinct early-life epigenomes that are enriched for epigenetic regulators and bona fide oncogenes. **A)** The heatmap reports z-score(log1p) transformed beta values of differentially methylated probes in *Trim28^+/D9^*-*heavy* vs -*light* animals. Effect size cut-off = 0.1; p-value cut-off = 0.001. N=24 (15 *Trim28^+/D9^*-*heavy* vs 9 *Trim28^+/D9^*-*light*). **B)** Enriched features (<u>upper panel</u>) and genes (<u>lower</u> <u>panel</u>) for differentially methylated probes between *Trim28^+/D9^*-*heavy* vs -*light* animals. False Discovery Rate (FDR) cut-off = 0.01 (one-tailed Fisher’s exact test). **C)** Gene-ontology (GO) enrichment plot of biological processes enriched in DML between *Trim28^+/D9^*-*heavy* vs -*light* animals. The size of the dots represents the gene-ratio, while the color represents the adjusted p-value for each term; Benjamini–Hochberg adjusted p-value cut-off = 0.05. **D)** Odds ratio from Fisher’s exact tests, testing the statistical significance of overlaps between genes enriched by hypo-methylated probes in *Trim28^+/D9^*-*heavy* (dark orange) and -*light* animals (pale orange), and the COSMIC Cancer Gene Census (https://cancer.sanger.ac.uk/census). P-value = 0.0002, *** p<0.001. **E)** <u>Left panel</u>: Kaplan-Meier survival probability as percentage of the total population and the time of survival in months. Log-rank test, p=1.516e^-6^. Total cases analyzed = 10967. <u>Right</u> <u>panel</u>: Kaplan-Meier disease-free survival probability. Log-rank test, p=1.27e^-9^. In both panels all TCGA PanCancer Atlas patients mutated in genes from the *Trim28^+/D9^*-*light* hypo-methylated signature (pale orange, n=3766) are compared to individuals not mutated in the same genes (black, n=7180). **F)** Heatmap of the effects on overall survival probability of mutations in the indicated genes and tumor tissues. The analysis includes all samples from TCGA and non-TCGA studies with no overlapping samples, from cBioPortal (N=69223 samples). Tumor tissues are separated in two main branches according to sample number informing the analysis (left: >3000 samples; right: <3000 samples). **G)** Volcano plot showing the type of interaction for pairwise mutations in genes from the *Trim28^+/D9^*-*light* hypo-methylated signature, in all TCGA PanCancer Atlas patients. Red indicates co-occurrence and blue mutually exclusivity of pairwise mutations. One-sided Fisher’s Exact Test; Benjamini–Hochberg adjusted p-value cut-off = 0.05. Total cases analyzed = 10967. **H)** Our model suggests that TRIM28 buffers intrinsic developmental heterogeneity via heterochromatin silencing. By modulating a differentially methylated cancer-related gene set primes two distinct developmental trajectories for cancer susceptibility and outcomes, with one of the trajectories being more resistant and the other prone to cancer.

Enrichments were also observed for probes within the *epigenetic aging clock*; *enhancers*; and at select subsets of transcriptional start sites (*TssBiv* and *TssFlnk*; **Fig 3B** top panel, green and red). Collectively, these data suggest that *Trim28^+/D9^-light* animals have more permissive chromatin at regions that would otherwise be silenced. To our knowledge, these data represent the first epigenetic signatures of developmentally programmed cancer susceptibility states.

### The *Trim28^+/D9^*-sensitive epigenome is enriched for epigenetic regulators and bona fide oncogenes

We next examined the specific *genes* that were differentially methylated between the -*light* and -*heavy* morphs at 10 days of age (DMGs; **Fig. 3B** bottom panel). Interestingly, DMGs hypomethylated in *Trim28^+/D9^-light* animals were also enriched in epigenetic regulators of *gene expression*, *heterochromatin formation*, *heterochromatin organization*, *genomic imprinting*, *DNA methylation* and *DNA alkylation,* among others (**Fig. 3B**, bottom panel, pale orange, and 3**C**). Therefore, *light*-morphs exhibit hypomethylation at coding regions of epigenetic silencers *and* their targeted genomic regions (**Fig. 3B**, top left panel).

We also queried three independent resources to gain insight into whether dysregulation of the -*light* and - *heavy* DMGs alone would be expected to impact cancer outcomes. We search for their enrichment in the Jensen DISEASE database of disease-gene associations^60^, and found that -*light* and -*heavy* DMGs were enriched for pathways underpinning human development and cancer (**Fig. S3J**), findings consistent with phenotypes in the mouse. Second, we tested for presence of -*light* / -*heavy* DMG orthologs in the COSMIC Cancer Gene Census (COSMIC), a database of high confidence oncogenes and tumor-suppressor genes^61^. In agreement with the *aggressive* MCS phenotype of -*light* morphs, DMGs ‘activated’ (hypomethylated) in *Trim28^+/D9^-light* animals were significantly enriched for known oncogenes (*GNAS*, *JAK3*, *MYCN*, *HMGA2*) (**Fig. 3D**). Finally, we used the TCGA PanCancer Atlas^62^ and compared cancer outcomes of individuals with or without mutations in *-light* or *-heavy* specific DMGs. Interestingly, patients with mutations in *Trim28^+/D9^-light* hypomethylated DMGs showed reduced overall survival probability (**Fig. 3E**, left panel) and a striking difference in time-to-relapse (**Fig. 3E**, right panel) compared to patients bearing other mutations. Stratifying the data by tumor-type showed wide-spread tumor promoting effects in orthologs of essentially all *Trim28^+/D9^-light* DMGs, with significantly reduced survival rates across many tumor types (**Fig. 3F** and **S3K**). Noteworthy in the stratification analysis, prostatic adenocarcinoma showed cancer-accelerating associations with orthologs of almost all -*light* specific DMGs (**Fig. 3F** and **S3K**, leftmost column); prostatic adenocarcinoma was the most prevalent cancer type specifically in the *Trim28^+/D9^-light* animals (**Fig. 2G**). No differences were found for the same analyses performed for orthologs of hypermethylated DMGs (i.e., those hypomethylated in -*heavy* animals) (**Fig. S3L**). Collectively, these data identify the hypomethylated *Trim28^+/D9^-light* DMGs as putative mediators of the altered cancer susceptibility states found in *Trim28^+/D9^* animals.

As a final assessment of on-coregulatory potential, we tested for mutational co-occurrence of the same DMG ortholog sets in the TCGA PanCancer Atlas. Co-occurrence of mutations within groups of genes may suggest that those alleles can provide additive or synergistic tumor survival advantages. Consistent with the phenotype of the *Trim28^+/D9^-light* morphs above, co-occurrence of mutations in *-light* DMG orthologs was markedly over-represented in human primary tumors, suggesting that they are part of a co-regulated signature (**Fig. 3G**). Thus, the early-life *Trim28^+/D9^*-dependent epigenome is enriched for bona fide oncogenes. Collectively, these data suggest a model whereby intrinsic differences in early-life epigenetic programming may determine cancer outcomes (**Fig. 3H**).

## DISCUSSION

### Early-life epigenetic heterogeneity as a regulator of differential cancer susceptibility

Here we identify (TRIM28-buffered) intrinsic developmental heterogeneity as a novel determinant of cancer susceptibility. The data show that TRIM28 haploinsufficiency generates two reproducible developmental morphs at the organismal level and that these differ in their cancer susceptibility; one “resistant”, and one more “susceptible”. Conceptually, the result is similar to epigenetic heterogeneity described within tumors^63^ and between tumors^64,65^, except at the inter-organismal level. How meta-stable states between identical cells or organisms are imposed remains unclear, though pioneering work on meta-stability of variegating reporters implicate epigenetic silencing machinery^66–70^. Along those lines, it has been suggested that a key condition for the emergence of alternate *cellular* states is the epigenetic reorganization of the genome^34,71,72^. Feinberg and Levchenko^34^ recently provided an innovative theoretical framework for how genetic and epigenetic networks generate meta-stable partitions and alternate cellular functional states, a potential energy landscape model that include energy wells or ‘attractors’. DNA mutations and/or changes in epigenetic topology (via DNA mutation, DNA methylation, or histone modifications, for instance) alter that landscape and create new/alternate attractor states. Our data suggest that these same concepts hold true at the *organismal* scale, and that these differences can have real consequences for cancer outcomes. They suggest that TRIM28-dependent silencing helps define the shape of the potential energy landscape (e.g., by controlling the depth of or barrier between attractor states). In the same way that oncogenic mutations have different effects depending on cellular developmental stage^73–77^, our data suggest that oncogenesis can also be influenced by stochastic, organismal epigenetic programs that are established in development.

### Role of epigenetics in the developmental origin of cancer

This study is one of the first attempts to bridge two key questions in the field of cancer epigenetics. One is a “cell of origin” question: how does the epigenetic state of a cell permit, support, or resist oncogenic transformation^14,16–23^? The second is: how do early-life epigenetic cues change or modulate cancer risk (between individuals)^35,36,78,79^? Addressing these questions requires isogenic models and identical environments, in part because individual development involves stochastic processes.

Our data show that inter-individual differences in early-life epigenome organization can dictate differential cancer development, prevalence, and survival. This finding complements prior work indicating that H3K9me3 (a target of TRIM28 action) most strongly correlates with tumor mutation density^20^. Together with the latter work, the inter-individual epigenetic differences identified in our current study suggest that one potential mechanism for the observed differential cancer outcomes is an altered sensitivity to mutations between the two morphs. We also detected important differences in Polycomb-targeted genes, poised, and bivalent regions between the cancer susceptibility morphs. During tumorigenesis, these regions are particularly sensitive to regulation via DNA methylation and may be correlated with cell “stemness”^80–83^. Regardless, the provocative implication arising from our data is that individual cancer susceptibility may have just as much to do with the epigenetic ‘background’ we are born with as it does DNA mutations, external environmental insults, or the cell of origin. Just as prior work demonstrated that epigenetic dysregulation at specific genes drives tumorigenesis in specific tissues or developmental stages, we would expect that the DMLs and DMGs identified here also have tissue-specific and developmental-stage specific effects. Analogous situations arise when different tumors exhibit mutations in different genes from a common or shared biochemical pathway^84–92^, or when cancer-associated mutations have different effects depending on the cell differentiation stage^93–97^. It would be interesting to understand when (during development) the different epigenetic backgrounds in the *Trim28^+/D9^* cancer susceptibility morphs become “activated”, and why some tissues seem to be more sensitive to tumorigenesis than others.

These novel findings differ from TRIM28’s published roles^42,98^ that include context-specific oncogene^99–109^ and tumor suppressor^110–115^ functions, roles that were primarily determined from complete (homozygous) knockouts. In contrast to homozygous knockout models, the *Trim28^+/D9^* haplo-insufficient mouse exhibits near normal levels of TRIM28 throughout the body^44^. Given TRIM28’s presence in multiple complexes (Trim28/KAP1 co-repressor complex, NuRD, CoREST, PML-NB, BORG/TRIM28, ZMYM2-TRIM28, MAGE-Trim28, or HUSH), this is an important distinction between the models. Indeed, TRIM28 homozygous deletion is embryonic lethal^116–118^. The DNA-methylation differences between -*light* and -*heavy* morphs suggest that *Trim28^+/D9^* specifically impacts TRIM28’s silencing function. This fundamental difference between models may explain for instance why TRIM28 knockout models develop liver tumors^111,112^, and *Trim28^+/D9^* mice do not.

### Limitations of the study and future directions

This work provides proof-of-concept that early-life, epigenetically distinct developmental programs can result in differential cancer susceptibility. The ability to show this effect in multiple tissue types is both a strength and a limitation of this study. Indeed, by focusing on an MCS model, we demonstrate that differential susceptibility is a property of the entire organism and can identify responsive tissues. At the same time, thousands of animals would be needed to draw the same conclusions for all the observed cancer sub-types including rare cancers. Likewise, and because the *Trim28^+/D9^* mutation in this model occurs in the whole body, it will be difficult to use the *Trim28^+/D9^* mouse (by itself) to dissect the molecular mechanisms down-stream of TRIM28 and heterochromatin disruption that underlies each of the observed cancers. A natural extension of this study is therefore to refine the model to understand the mechanistic basis of developmentally ‘primed’ cancer susceptibility for each cancer type.

Our data show that the epigenetic distinction between the two cancer susceptibility morphs is already evident by day 10, before weaning. Other open questions therefore relate to the precise timing of the observed (epigenetic) bifurcation, and the nature of any cell-intrinsic factors that might skew development towards one or the other epigenetic background. Our finding that the *Trim28^+/D9^* DMLs and DMGs are enriched for known human oncogenes and that they suggest a more permissive chromatin state at otherwise silenced regions, hints to a possible generalization of the model. If we can identify similarly sensitive regions of the human cancer genome, then we will be better equipped to optimally stratify and treat patients.

## METHODS

### Origin and maintenance of mice

*FVB/NJ.Trim28^+/MommeD9^* animals (*Trim28^+/D9^*) were generated in the Whitelaw lab^119^. *B6.129S4-Trp53<tm3.1Tyj>/J* animals (*Tp53^+/R270H^*) were originally generated in the Jacks lab^45^ and purchased from Jackson Laboratories (stock #008182). Both lines were fully backcrossed for more than 10 generations (FVB/NJ and B6 respectively). After bringing lines in-house, they were both maintained by internal breeding with wild-type siblings. Approximately 270 F1 hybrids were generated by crossing 8-week-old *FVB.Trim28^+/D9^* males with 8-week-old *B6.Tp53*^R270H/+^ females. One male was crossed with two females in the same cage, and females separated after plug checking the next morning. From these crosses, we generated 137 males (18 WT, 44 *Tp53^+/R270H^*, 34 *Trim28^+/D9^*, and 41 *Tp53^R270H/+^;Trim28^+/D9^*) and 133 females (30 WT, 36 *Tp53^+/R270H^*, 32 *Trim28^+/D9^*, and 35 *Tp53^R270H/+^;Trim28^+/D9^*). 114 animals were screened for tumors: 60 males (6 WT, 17 *Tp53^+/R270H^*, 15 *Trim28^+/D9^*, and 22 *Tp53^R270H/+^;Trim28^+/D9^*) and 54 females (8 WT, 21 *Tp53^+/R270H^*, 8 *Trim28^+/D9^*, and 17 *Tp53^R270H/+^;Trim28^+/D9^*). Only male offspring were analyzed; females exhibited unusually low levels of both phenotypic heterogeneity and cancer incidence precluding analysis.

All animals were fed breeder chow (Lab diet, 5021 cat. #0006540) *ad libitum* and housed in individually ventilated cages (Tecniplast, Sealsafe Plus GM500 in DGM Racks). All animals were kept on a 12-hour light/dark cycle at an average ambient temperature of 23 °C and 35% humidity. The maximum capacity per cage is 5 animals, and each cage was enriched with Enviro-dri (The Andersons, Crink-l’Nest) and cardboard dome homes (Shepherd, Shepherd Shack Dome). Whenever possible, same-sex siblings and same-sex animals from different litters were combined (∼20 days of age) to co-house isogenic animals. At 4, 8, 16, 32, 40, 50, 60, and 70 weeks of age (or at euthanasia), mice were weighed and scanned via EchoMRI for fat and lean mass composition (EchoMRI™, EchoMRI™-100H), in the morning. All protocols were approved by Institutional Animal Care and Use Committee under protocols 19-0026, 22-09-036, 18-10-028, and 21-08-023, at Van Andel Institute (VAI, USA).

### Genotyping

Ear punch biopsies were collected at 10 days and placed in a 20 µl reaction mix composed of genomic DNA lysis buffer (100 mM Tris-HCl pH 8.5, 5 mM EDTA, 0.2% SDS, 100 mM NaCl) supplemented with 20 mg proteinase K (Thermo Scientific, EO0491). Biopsies were digested with a thermal cycling protocol consisting of 55 °C for 16 hours, 95 °C for 10 minutes, and a 4 °C hold (lid at 105 °C). Thereafter, 160 µl of nuclease-free water (Invitrogen, AM9938) was added to each lysate to achieve 180 µl final volume. The PCR reaction (Thermo Scientific, EP0703) for *Trim28* and *Tp53* alleles comprised 1 µl diluted biopsy lysate and 19 µl reaction master mix (1X DreamTaq Buffer, 0.2 mM dNTPs, 0.1 µM primer forward and reverse mix, 2 U DreamTaq DNA Polymerase, in nuclease-free water), with PCR primer and thermal cycling conditions listed in the Supplementary Tables 1 and 2, respectively. To verify the presence of each point-mutation, 20 µl of each PCR product was digested with either 0.5 µl XceI/NspI (for *Trim28^+/D9^*; Themo Scientific, FD1474) or 0.5 µl MslI (for *Tp53^R270H^*^/+^; New England BioLabs, R0571L) in a final reaction volume of 30 µl. Digestion products (∼700 bp WT *Trim28*, ∼250 bp + ∼450 bp *Trim28^+/D9^*, ∼500 bp WT *Tp53*, ∼200 + ∼300 bp *Tp53^R270H/+^*) were detected on a 3% agarose gel (Fisher Scientific, BP160-500) in 1X TAE, with GelRed as the intercalating dye (Biotium, 41003).

### Statistical analyses of developmental heterogeneity

We used Levene’s test^120^ on body, fat, or lean masses independently to test for homoscedasticity (or equality of variances) across genotypes. P-values were subsequently adjusted for multiple comparisons using the Benjamini-Hochberg method^121^. We used MClust (version 5.4.9)^122^ for iterative Expectation-Maximization (EM) maximum-likelihood estimation in parameterized Gaussian mixture models. We chose to regularize with a prior to achieve smoother Bayesian Information Criterion (BIC). The uncertainty in the classification was used as graphical parameter for the lean/fat mass data plots, and for weighing the Log-Rank p-values in mouse Kaplan-Meier plots. Since most mice were classified with high confidence, the effect of this correction is negligible. We validated the results from MClust with Rmixmod (version 2.1.8)^123^ using unsupervised classification and density estimation with 3 criteria: BIC, ICL, and NEC. Both methods were used to cluster 16-week fat and lean mass data for each genotype.

We used generalized additive models (GAMs)^124^ to model the fat or lean mass changes over time per group. We used random-effect splines to model a random slope and random intercept for each animal (by week). We then used the “emmeans” R package^125,126^ to compare the overall slope of the fat or lean mass by group. We ran a separate GAM to model the differences in fat or lean mass within each timepoint. For this model, we included a random-effect spline for each animal but excluded the spine for a random slope by week. Again, we used the “emmeans” package to compare the differences in the fat or lean mass within each timepoint. We used a two-sample test of proportions^127^ under the “stats” R package^128,129^ to examine the differences in proportion of animals that died with cancer before and after the endpoint in the *Trim28 -heavy* and *-light* trajectories. The p-values from those tests were adjusted for multiple testing using the Benjamini-Hochberg method.

### Health monitoring

Professional vivarium staff checked mice for overall health, well-being, and the presence of any abnormal mass/tumor 2-3 times per week. Mice were euthanized if they showed any of the following symptoms: >20% weight loss, tumors that were ∼15% of total body weight, tumor ulcerations, discharge or hemorrhage from the tumor, limited ambulation, reduced appetite and drinking, limited defecation or urination, abnormal gait or posture, labored breathing, lack of movement, or hypothermia. Mice with reported health issues or that reached the study endpoint (70 weeks of age) were euthanized via CO_2_ asphyxiation and cervical dislocation.

### Tissue harvesting

The following tissues were dissected and immediately fixed in 10% NBF solution (3.7-4% formaldehyde 37-40%, 0.03 M NaH_2_PO_4_, 0.05 M Na_2_HPO_4_, in distilled water with final pH of 7.2± 0.5): epidydimal white adipose tissue (eWAT); uterus or preputial glands, seminal vesicles, and testis (depending on sex); bladder; pancreas; spleen; intestine; stomach; mesenteric fat; liver; kidneys; heart; lungs; thymus; brain; the ninth breast; hindlimb muscles; and hindlimb bones. We also recovered spine, ribs, skull, skin, and any other mass if they appeared to be involved with a tumor or disease. The volume of fixative was at least 15-20 times greater than the volume of tissue. Specimens > 2.5 mm thick were cut to ensure adequate fixation. All the tissues but eWAT, mesenteric fat, uterus, and bones (including spine) were fixed for 48 hours. The fat-rich tissues (eWAT, mesenteric fat, uterus) were fixed for 72 hours. The bony tissues (bones and spine) were fixed for 1 week followed by 1 week decalcification in 14% EDTA (14% free-acid EDTA at pH 7.2, adjusted with NH₄OH). After each incubation, all the tissues were moved to 70% ethanol.

### Tissue preparation for histology

All tissues were embedded in paraffin by the Van Andel Institute Pathology and Biorepository Core following internal standard operating procedures. Upon receipt, tissues were dehydrated and cleared using a Tissue-Tek VIP 5 (Sakura) and an automated protocol consisting of 60’ in 70% ethanol; 60’ in 80% ethanol; 2 x 60’ in 95% ethanol; 3 x 60’ in 100% ethanol; 2 x 30’ in xylene; and 1 x 30’ and 1 x 45’in paraffin. Tissues were embedded in paraffin with a Leica EG1150 embedding center. Three 5-µm thin sections spaced 150 µm apart were cut from each tissue for hematoxylin and eosin (H&E) staining using a Leica rotary microtome. The remaining tissue was conserved as a paraffin embedded tissue block. H&E staining was performed with a Tissue-Tek Prisma Plus Automated Slide Stainer (Sakura) and Tissue-Tek Prisma H&E Staining Kit #1.

### Pathology evaluation

Standard 5-µm thick tissue sections stained with H&E were assessed for tumors and dysplastic lesions by a board-certified pathologist dedicated to this study. Most samples were provided to the pathologist in a blinded manner. Tumors were broadly classified as carcinomas, germ cell tumors, leukemias, lymphomas, and sarcomas. A detailed classification was provided based on the tissue of origin.

### Mouse DNA methylation array

Ear punch biopsies were collected as described above, and DNA purified from lysate using a DNeasy Blood & Tissue Kit (QIAGEN, 69504) with slight modifications. After tissue digestion, the lysate was brought to 220 µl with 1X PBS. We then we followed steps 2-7 of the Quick-Start protocol. DNA was eluted with two washes of 100 µl Buffer AE, and purified DNA quantified by Qubit fluorometry (Life Technologies). Then, 6-500 ng of DNA from each sample was bisulfite-converted using the Zymo EZ DNA Methylation Kit (Zymo Research, Irvine, CA USA) following the manufacturer’s protocol and the specified modifications for the Illumina Infinium methylation assays. After conversion, all bisulfite reactions were cleaned using the Zymo-Spin binding columns and eluted in 12 µL of Tris buffer.

Following elution, bisulfite-converted and restored DNA was processed through the Illumina mouse methylation array protocol. The mouse methylation array contains >285K probes for CpG islands, translation start sites, enhancers, imprinted loci, and other regions, along with strain-specific SNP probes^130^. To perform the assay, 7 µl of converted DNA was denatured with 4 µl 0.1N sodium hydroxide. DNA was then amplified, hybridized to the Infinium bead chip, and an extension reaction performed using fluorophore-labeled nucleotides per the manufacturer’s protocol. Arrays were scanned on the Illumina iScan platform, and probe-specific calls were made using Illumina Genome Studio v2011.1 software to generate IDAT files.

### DNA methylation analysis

Analysis of IDAT files was performed using the default SeSAMe pipeline^131^ and its wrapper pipeline SeSAMeStr^132^. Fifty-eight independent biological replicates from WT, *Trim28*, *Tp53* and *Trim28/Tp53* male animals were analyzed. Data pre-processing and quality controls were performed using SeSAMe default parameters and the pre-processing code ‘TQCDPB’. All samples showed a detection rate >90% and no dye bias. In all differential DNA methylation analyses, the effect size cutoff was set to 0.1 (i.e., 10% differential DNA methylation) and the p-value cutoff was <0.05, unless otherwise specified in the figure legends. For all differential DNA methylation analyses between isogenic *Trim28* - *heavy* and *-light* animals, the effect size cutoff was set to 0.05 (i.e., 5% differential DNA methylation) and the p-value cutoff was <0.05. For all differential analyses, we included batch effect as a covariate in the model. Other technical and biological effects/bias (i.e., detection rate, initial DNA concentration, litter) were evaluated but not included in the model because they were co-linear with the batch effect and/or did not separate groups in dimensional reduction analyses. Global DNA methylation correlation analysis was performed using the ‘chart.Correlation’ function from the ‘PerformanceAnalytics’ R package. Similarity between samples was calculated as the sum of squared residuals from linear regressions. Principal component analysis (PCA) of beta values was performed on SeSAMeStr pipeline output using the R function ‘prcomp’. Heatmap visualization of differentially methylated loci (DML) was performed using the R package ‘ComplexHeatmap’^133^. For heatmap visualization, beta values were modelled and weighted using the Mclust certainty score in limma^134^. Probe enrichment analysis was performed using SeSAMe KnowYourCG module. Gene ontology analysis of probes-enriched genes was performed using the R package ‘clusterProfiler’^135^. Gene enrichment in the Jensen DISEASES database^136^ was performed using the R package ‘enrichR’^137^. Further data visualization of SeSAMe/SeSAMeStr output was perform in R using Rstudio.

### TCGA PanCancer Atlas data analyses

The TCGA PanCancer Atlas^62^ encompasses 32 studies and 10967 samples. All preliminary analyses were performed using cBioPortal^138,139^, and outputs were used to replot and visualize the data in Rstudio. All Kaplan-Meier curves were generated by selecting all TCGA PanCancer Atlas cases that harbor mutations in either the *Trim28^+/D9^ -heavy* or *-light* gene signatures, and comparing them with cases without mutations in the same genes. Statistical significance for differences between groups in all Kaplan-Meier curves was tested by log-rank tests with a p-value cut-off = 0.05. Mutual co-occurrence or exclusivity of pairwise mutations in genes within either the *Trim28^+/D9^ -heavy* or *-light* gene signatures were tested by one-sided Fisher’s Exact Test, with a Benjamini–Hochberg adjusted p-value cut-off = 0.05. Statistical significance of overlaps between genes within either the *Trim28^+/D9^ -heavy* or *-light* gene signatures and the COSMIC Cancer Gene Census^61^ was tested by a Fisher’s exact test with a p-value cut-off = 0.05. The effect of mutations at the *Trim28^+/D9^ -light* signature genes was assessed in each group of tumors from all available tissue types in cBioPortal. This analysis comprised all samples from TCGA and non-overlapping samples from cBioPortal (N=69223 samples). Definition of ‘harmful’ and ‘protective’ mutations was based on the ratio of the median months survival between the samples with no mutations and the samples carrying mutations (i.e., median months survival unaltered/altered samples >1 = ‘harmful’ mutation; median months survival unaltered/altered samples <1 = ‘protective’ mutation). As the number of significant hits in this analysis is biased by the total number of samples for each targeted tissue, we split the tumor tissues into low (<3000) and high (>3000) samples number and showed them separated in the heatmap visualization. The same analysis was performed on the overall survival and the disease-free survival.

## Data and code availability

All DNA methylation array data generated in this study were deposited to Gene Expression Omnibus (GEO) under accession code GSE229030 The SeSAMe wrapper pipeline SeSAMeStr is published online in Zenodo: https://doi.org/10.5281/zenodo.7510575. No other custom code or mathematical algorithms were generated for this study. All publicly available codes and tools used to analyze the data are reported and referenced in the methods sections. Any additional information required to reanalyze the data reported in this paper is available from the lead contact upon request.

## ACKNOWLEDGEMENTS

We thank P. Laird, H. Shen, T. Yang, R. Jones, B, Williams, E. Lien, C. Essenburg, N. Vander Schaaf, P. Stevens, L. DeCamp, E. Levine, E. Ma, D. Lu, H. Lu, V. Molchanov, J. Endicott, V. Wegert, M. Edozie, and D. Aicher for technical insight, suggestions, and support. This work would not have been possible without the amazing support of MPI-IE Facilities and VAI Vivarium (in particular, B. Eagleson, S. Bechaz, A. Rapp, R. Burdette, E. Tubbergen, E. Hamel, M. Powers, and S. Greenwald; RRID:SCR_023211); Transgenics (in particular, T. Kempston, and A. Guikema; RRID:SCR_022914); Pathology and Biorepository (in particular, S. Jewell, D. Rohrer, B. Berghuis, L. Turner, S. Whitford, A. Bouwman, K. Feenstra, and K. Goudreau; RRID:SCR_022912); Bioinformatics and Biostatistics, and Genomics (in particular, M. Adams, M. Wegener, and T. Avequin; RRID:SCR_022913) Cores. We thank P. Laird, E. Lien, and J. Jang for critical evaluation of the manuscript. This work was supported by funding from the MPG, the ERC, Van Andel Institute through internal philanthropy and MeNU pilot project grants, NIH award number 1R01HG012444 (to AP), R01AI171984 and Chan Zuckerberg Initiative with award number DI-000000287 (To TJTJ), and Human Frontier Science Program Long-Term Fellowship LT000441/2018-L (to IP).

## AUTHOR CONTRIBUTIONS

IP and JAP conceived the project. IP, JAP, DS, and TJTJ designed the overall methodology and IP designed each individual experiment. IP and MT maintained and performed the *in vivo* experiments, genotyping and most of the tissue harvesting. IP performed DNA extraction for methylation analysis. YL, KS, YC, AB, AD, ED, and members of the PERMUTE group supported the *in vivo* experiments and the genotyping. LF and SA developed the SeSAMeStr package and curated the data. IP analyzed all the phenotypic data, while LF and SA performed methylation data analysis. GH performed all the pathology reviews. IP, EW, ZM, and TJTJ performed the statistical analyses of developmental heterogeneity. IP, LF, DC and JAP wrote the original draft. IP and LF prepared the figures. IP, LF, DC, JAP, DS reviewed and edited the initial draft. JAP, TJTJ, DC and IP acquired funds. JAP provided resources for the experiments. JAP, TJTJ and IP supervised the work.

## CONSORTIA

### PERMUTE

J. Andrew Pospisilik, Ilaria Panzeri, Luca Fagnocchi, Stefanos Apostle, Emily Wolfrum, Zachary Madaj, Jillian Richards, Holly Dykstra, Tim Gruber, Mitch McDonald, Andrea Parham, Brooke Armistead, Timothy J. Triche Jr., Zachary DeBruine, Mao Ding, Ember Tokarski, Eve Gardner, Joseph Nadeau, Christine Lary, Carmen Khoo, Ildiko Polyak, Qingchu Jin.

## DECLARATION OF INTERESTS

The authors declare no competing interests.

## Notes

### Competing Interest Statement

The authors have declared no competing interest.

## REFERENCES

1 Blanpain, C. Tracing the cellular origin of cancer. Nat. Cell Biol. 15, 126–134, doi:10.1038/ncb2657 (2013).

2 Merlo, L. M., Pepper, J. W., Reid, B. J. & Maley, C. C. Cancer as an evolutionary and ecological process. Nat. Rev. Cancer 6, 924–935 (2006).

3 Greaves, M. & Maley, C. C. Clonal evolution in cancer. Nature 481, 306–313, doi:10.1038/nature10762 (2012).

4 Martincorena, I. et al. High burden and pervasive positive selection of somatic mutations in normal human skin. Science (New York, N.Y.) 348, 880–886, doi:doi:10.1126/science.aaa6806 (2015).

5 Yizhak, K. et al. RNA sequence analysis reveals macroscopic somatic clonal expansion across normal tissues. Science (New York, N.Y.) 364, doi:10.1126/science.aaw0726 (2019).

6 García-Nieto, P. E., Morrison, A. J. & Fraser, H. B. The somatic mutation landscape of the human body. Genome Biol. 20, 298, doi:10.1186/s13059-019-1919-5 (2019).

7 Bose, S., Deininger, M., Gora-Tybor, J., Goldman, J. M. & Melo, J. V. The presence of typical and atypical BCR-ABL fusion genes in leukocytes of normal individuals: biologic significance and implications for the assessment of minimal residual disease. Blood 92, 3362–3367 (1998).

8 Lodato, M. A. et al. Aging and neurodegeneration are associated with increased mutations in single human neurons. Science (New York, N.Y.) 359, 555–559, doi:10.1126/science.aao4426 (2018).

9 Lee-Six, H. et al. The landscape of somatic mutation in normal colorectal epithelial cells. Nature 574, 532–537, doi:10.1038/s41586-019-1672-7 (2019).

10 Ganz, J. et al. Rates and patterns of clonal oncogenic mutations in the normal human brain. Cancer Discov. 12, 172–185, doi:10.1158/2159-8290.Cd-21-0245 (2022).

11 Acha-Sagredo, A., Ganguli, P. & Ciccarelli, F. D. Somatic variation in normal tissues: friend or foe of cancer early detection? Ann. Oncol. 33, 1239–1249, doi:10.1016/j.annonc.2022.09.156 (2022).

12 Kakiuchi, N. & Ogawa, S. Clonal expansion in non-cancer tissues. Nat. Rev. Cancer 21, 239–256, doi:10.1038/s41568-021-00335-3 (2021).

13 Jassim, A., Rahrmann, E. P., Simons, B. D. & Gilbertson, R. J. Cancers make their own luck: theories of cancer origins. Nature Reviews Cancer, doi:10.1038/s41568-023-00602-5 (2023).

14 Schneider, G., Schmidt-Supprian, M., Rad, R. & Saur, D. Tissue-specific tumorigenesis: context matters. Nature Reviews Cancer 17, 239–253, doi:10.1038/nrc.2017.5 (2017).

15 Haigis, K. M., Cichowski, K. & Elledge, S. J. Tissue-specificity in cancer: The rule, not the exception. Science (New York, N.Y.) 363, 1150–1151, doi:doi:10.1126/science.aaw3472 (2019).

16 Baggiolini, A. et al. Developmental chromatin programs determine oncogenic competence in melanoma. Science (New York, N.Y.) 373, eabc1048, doi:doi:10.1126/science.abc1048 (2021).

17 Berquam-Vrieze, K. E. et al. Cell of origin strongly influences genetic selection in a mouse model of T-ALL. Blood 118, 4646–4656, doi:10.1182/blood-2011-03-343947 (2011).

18 Hinoue, T. et al. Analysis of the association between CIMP and BRAFV600E in colorectal cancer by DNA methylation profiling. PloS one 4, e8357 (2009).

19 Polak, P. et al. Cell-of-origin chromatin organization shapes the mutational landscape of cancer. Nature 518, 360–364, doi:10.1038/nature14221 (2015).

20 Schuster-Böckler, B. & Lehner, B. Chromatin organization is a major influence on regional mutation rates in human cancer cells. Nature 488, 504–507, doi:10.1038/nature11273 (2012).

21 Vicente-Dueñas, C., Hauer, J., Cobaleda, C., Borkhardt, A. & Sánchez-García, I. Epigenetic Priming in Cancer Initiation. Trends Cancer 4, 408–417, doi:10.1016/j.trecan.2018.04.007 (2018).

22 Visvader, J. E. Cells of origin in cancer. Nature 469, 314–322, doi:10.1038/nature09781 (2011).

23 Yamamoto, E. et al. Molecular Dissection of Premalignant Colorectal Lesions Reveals Early Onset of the CpG Island Methylator Phenotype. The American Journal of Pathology 181, 1847–1861, 10.1016/j.ajpath.2012.08.007 (2012).

24 Castillo-Fernandez, J. E., Spector, T. D. & Bell, J. T. Epigenetics of discordant monozygotic twins: implications for disease. Genome Medicine 6, 60, doi:10.1186/s13073-014-0060-z (2014).

25 Angers, B., Perez, M., Menicucci, T. & Leung, C. Sources of epigenetic variation and their applications in natural populations. Evolutionary Applications 13, 1262–1278, 10.1111/eva.12946 (2020).

26 Machin, G. Non-identical monozygotic twins, intermediate twin types, zygosity testing, and the non-random nature of monozygotic twinning: a review. Am J Med Genet C Semin Med Genet 151c, 110–127, doi:10.1002/ajmg.c.30212 (2009).

27 Youssoufian, H. & Pyeritz, R. E. Mechanisms and consequences of somatic mosaicism in humans. Nat Rev Genet 3, 748–758, doi:10.1038/nrg906 (2002).

28 de Oliveira Andrade, F., et al. Exposure to lard-based high-fat diet during fetal and lactation periods modifies breast cancer susceptibility in adulthood in rats. The Journal of nutritional biochemistry 25, 613–622, doi:10.1016/j.jnutbio.2014.02.002 (2014).

29 Ekbom, A., Adami, H. O., Trichopoulos, D., Hsieh, C. C. & Lan, S. J. Evidence of prenatal influences on breast cancer risk. The Lancet 340, 1015–1018, 10.1016/0140-6736(92)93019-J (1992).

30 Murugan, S., Zhang, C., Mojtahedzadeh, S. & Sarkar, D. K. Alcohol exposure in utero increases susceptibility to prostate tumorigenesis in rat offspring. Alcoholism, clinical and experimental research 37, 1901–1909, doi:10.1111/acer.12171 (2013).

31 Prins, G. S. Endocrine disruptors and prostate cancer risk. Endocrine-related cancer 15, 649–656, doi:10.1677/erc-08-0043 (2008).

32 Rinaldi, J. C. et al. Implications of intrauterine protein malnutrition on prostate growth, maturation and aging. Life sciences 92, 763–774, doi:10.1016/j.lfs.2013.02.007 (2013).

33 Sarkar, D. K. Fetal alcohol exposure increases susceptibility to carcinogenesis and promotes tumor progression in prostate gland. Advances in experimental medicine and biology 815, 389–402, doi:10.1007/978-3-319-09614-8_23 (2015).

34 Feinberg, A. P. & Levchenko, A. Epigenetics as a mediator of plasticity in cancer. Science (New York, N.Y.) 379, eaaw3835, doi:doi:10.1126/science.aaw3835 (2023).

35 Herceg, Z. et al. Roadmap for investigating epigenome deregulation and environmental origins of cancer. International journal of cancer 142, 874–882, doi:10.1002/ijc.31014 (2018).

36 Ho, S.-M. et al. in The Epigenome and Developmental Origins of Health and Disease (ed Cheryl S. Rosenfeld) 315–336 (Academic Press, 2016).

37 Ivanov, A. V. et al. PHD Domain-Mediated E3 Ligase Activity Directs Intramolecular Sumoylation of an Adjacent Bromodomain Required for Gene Silencing. Mol Cell 28, 823–837, doi:10.1016/j.molcel.2007.11.012 (2007).

38 Quenneville, S. et al. In embryonic stem cells, ZFP57/KAP1 recognize a methylated hexanucleotide to affect chromatin and DNA methylation of imprinting control regions. Mol Cell 44, 361–372, doi:10.1016/j.molcel.2011.08.032 (2011).

39 Ryan, R. F. et al. KAP-1 corepressor protein interacts and colocalizes with heterochromatic and euchromatic HP1 proteins: a potential role for Krüppel-associated box-zinc finger proteins in heterochromatin-mediated gene silencing. Mol Cell Biol 19, 4366–4378, doi:10.1128/MCB.19.6.4366 (1999).

40 Schultz, D. C., Ayyanathan, K., Negorev, D., Maul, G. G. & Rauscher, F. J. SETDB1: a novel KAP-1-associated histone H3, lysine 9-specific methyltransferase that contributes to HP1-mediated silencing of euchromatic genes by KRAB zinc-finger proteins. Genes & development 16, 919–932 (2002).

41 Schultz, D. C., Friedman, J. R. & Rauscher, F. J., 3rd. Targeting histone deacetylase complexes via KRAB-zinc finger proteins: the PHD and bromodomains of KAP-1 form a cooperative unit that recruits a novel isoform of the Mi-2alpha subunit of NuRD. Genes & development 15, 428–443, doi:10.1101/gad.869501 (2001).

42 Czerwińska, P., Mazurek, S. & Wiznerowicz, M. The complexity of TRIM28 contribution to cancer.Journal of Biomedical Science 24, 63, doi:10.1186/s12929-017-0374-4 (2017).

43 Whitelaw, N. C. et al. Reduced levels of two modifiers of epigenetic gene silencing, Dnmt3a and Trim28, cause increased phenotypic noise. Genome Biology 11, R111, doi:10.1186/gb-2010-11-11-r111 (2010).

44 Dalgaard, K. et al. Trim28 Haploinsufficiency Triggers Bi-stable Epigenetic Obesity. Cell 164, 353–364, doi:10.1016/j.cell.2015.12.025 (2016).

45 Olive, K. P. et al. Mutant p53 gain of function in two mouse models of Li-Fraumeni syndrome. Cell 119, 847–860, doi:10.1016/j.cell.2004.11.004 (2004).

46 Ashe, A. et al. A genome-wide screen for modifiers of transgene variegation identifies genes with critical roles in development. Genome Biology 9, R182, doi:10.1186/gb-2008-9-12-r182 (2008).

47 Blewitt Marnie, E., et al. An N-ethyl-N-nitrosourea screen for genes involved in variegation in the mouse. Proceedings of the National Academy of Sciences 102, 7629–7634, doi:10.1073/pnas.0409375102 (2005).

48 Harvey, M., McArthur, M. J., Montgomery Jr, C. A., Bradley, A. & Donehower, L. A. Genetic background alters the spectrum of tumors that develop in p53-deficient mice. FASEB J. 7, 938–943, 10.1096/fasebj.7.10.8344491 (1993).

49 Lang, G. A. et al. Gain of function of a p53 hot spot mutation in a mouse model of Li-Fraumeni syndrome. Cell 119, 861–872, doi:10.1016/j.cell.2004.11.006 (2004).

50 Bertozzi, T. M. & Ferguson-Smith, A. C. Metastable epialleles and their contribution to epigenetic inheritance in mammals. Seminars in cell & developmental biology 97, 93–105, doi:10.1016/j.semcdb.2019.08.002 (2020).

51 Van Baak, T. E. et al. Epigenetic supersimilarity of monozygotic twin pairs. Genome Biology 19, 2, doi:10.1186/s13059-017-1374-0 (2018).

52 Planterose Jiménez, B., et al. Equivalent DNA methylation variation between monozygotic co-twins and unrelated individuals reveals universal epigenetic inter-individual dissimilarity. Genome Biology 22, 18, doi:10.1186/s13059-020-02223-9 (2021).

53 Marttila, S. et al. Methylation status of VTRNA2-1/nc886 is stable across populations, monozygotic twin pairs and in majority of tissues. Epigenomics 14, 1105–1124, doi:10.2217/epi-2022-0228 (2022).

54 van Dongen, J. et al. Identical twins carry a persistent epigenetic signature of early genome programming. Nature Communications 12, 5618, doi:10.1038/s41467-021-25583-7 (2021).

55 Kaminsky, Z. A. et al. DNA methylation profiles in monozygotic and dizygotic twins. Nat Genet 41, 240–245, doi:10.1038/ng.286 (2009).

56 Zhou, W. et al. DNA Methylation Dynamics and Dysregulation Delineated by High-Throughput Profiling in the Mouse. bioRxiv, 2022.2003.2024.485667, doi:10.1101/2022.03.24.485667 (2022).

57 Tovy, A. et al. p53 is essential for DNA methylation homeostasis in naïve embryonic stem cells, and its loss promotes clonal heterogeneity. Genes Dev. 31, 959–972, doi:10.1101/gad.299198.117 (2017).

58 Filipczak, P. T. et al. p53-suppressed oncogene TET1 prevents cellular aging in lung cancer. Cancer Res. 79, 1758–1768, doi:10.1158/0008-5472.Can-18-1234 (2019).

59 Panatta, E. et al. Metabolic regulation by p53 prevents R-loop-associated genomic instability. Cell Reports 41, 111568, 10.1016/j.celrep.2022.111568 (2022).

60 Grissa, D., Junge, A., Oprea, T. I. & Jensen, L. J. Diseases 2.0: a weekly updated database of disease-gene associations from text mining and data integration. Database (Oxford) 2022, doi:10.1093/database/baac019 (2022).

61 Sondka, Z. et al. The COSMIC Cancer Gene Census: describing genetic dysfunction across all human cancers. Nat. Rev. Cancer 18, 696–705, doi:10.1038/s41568-018-0060-1 (2018).

62 Weinstein, J. N. et al. The Cancer Genome Atlas Pan-Cancer analysis project. Nat. Genet. 45, 1113–1120, doi:10.1038/ng.2764 (2013).

63 Li, Z., Seehawer, M. & Polyak, K. Untangling the web of intratumour heterogeneity. Nature Cell Biology 24, 1192–1201, doi:10.1038/s41556-022-00969-x (2022).

64 Gupta, Piyush B. et al. Stochastic State Transitions Give Rise to Phenotypic Equilibrium in Populations of Cancer Cells. Cell 146, 633–644, 10.1016/j.cell.2011.07.026 (2011).

65 Goyal, Y. et al. Diverse clonal fates emerge upon drug treatment of homogeneous cancer cells. Nature, doi:10.1038/s41586-023-06342-8 (2023).

66 Muller, H. J. Types of visible variations induced by X-rays in Drosophila. Journal of genetics 22, 299–334 (1930).

67 Girton, J. R. & Johansen, K. M. Chromatin structure and the regulation of gene expression: the lessons of PEV in Drosophila. Adv Genet 61, 1–43, doi:10.1016/s0065-2660(07)00001-6 (2008).

68 Allshire, R. C. & Ekwall, K. Epigenetic Regulation of Chromatin States in Schizosaccharomyces pombe. Cold Spring Harb Perspect Biol 7, a018770, doi:10.1101/cshperspect.a018770 (2015).

69 Rakyan, V. K. et al. Transgenerational inheritance of epigenetic states at the murine <i>Axin</i><sup><i>Fu</i></sup> allele occurs after maternal and paternal transmission. Proceedings of the National Academy of Sciences 100, 2538–2543, doi:doi:10.1073/pnas.0436776100 (2003).

70 Morgan, H. D., Sutherland, H. G. E., Martin, D. I. K. & Whitelaw, E. Epigenetic inheritance at the agouti locus in the mouse. Nature Genetics 23, 314–318, doi:10.1038/15490 (1999).

71 Pujadas, E. & Feinberg, Andrew P. Regulated Noise in the Epigenetic Landscape of Development and Disease. Cell 148, 1123–1131, 10.1016/j.cell.2012.02.045 (2012).

72 Feinberg, A. P. & Irizarry, R. A. Stochastic epigenetic variation as a driving force of development, evolutionary adaptation, and disease. Proceedings of the National Academy of Sciences 107, 1757–1764, doi:doi:10.1073/pnas.0906183107 (2010).

73 Schmidt, L. et al. Germline and somatic mutations in the tyrosine kinase domain of the MET proto-oncogene in papillary renal carcinomas. Nat. Genet. 16, 68–73 (1997).

74 Park, W. S. et al. Somatic mutations in the kinase domain of the Met/hepatocyte growth factor receptor gene in childhood hepatocellular carcinomas. Cancer Res. 59, 307–310 (1999).

75 Ma, P. C. et al. c-MET mutational analysis in small cell lung cancer: novel juxtamembrane domain mutations regulating cytoskeletal functions. Cancer Res. 63, 6272–6281 (2003).

76 McCoy, M. L., Mueller, C. R. & Roskelley, C. D. The role of the breast cancer susceptibility gene 1 (BRCA1) in sporadic epithelial ovarian cancer. Reprod. Biol. Endocrinol. 1, 1–5 (2003).

77 Beer, S. et al. Developmental context determines latency of MYC-induced tumorigenesis. PLoS Biol. 2, e332 (2004).

78 Alonso-Curbelo, D. et al. A gene–environment-induced epigenetic program initiates tumorigenesis. Nature 590, 642–648, doi:10.1038/s41586-020-03147-x (2021).

79 Carbone, M. et al. Tumour predisposition and cancer syndromes as models to study gene-environment interactions. Nat Rev Cancer 20, 533–549, doi:10.1038/s41568-020-0265-y (2020).

80 Ohm, J. E. et al. A stem cell–like chromatin pattern may predispose tumor suppressor genes to DNA hypermethylation and heritable silencing. Nature genetics 39, 237–242 (2007).

81 Schlesinger, Y. et al. Polycomb-mediated methylation on Lys27 of histone H3 pre-marks genes for de novo methylation in cancer. Nature genetics 39, 232–236 (2007).

82 Teschendorff, A. E. et al. Age-dependent DNA methylation of genes that are suppressed in stem cells is a hallmark of cancer. Genome Res 20, 440–446 (2010).

83 Widschwendter, M. et al. Epigenetic stem cell signature in cancer. Nature genetics 39, 157–158 (2007).

84 Ilyas, M., Tomlinson, I., Rowan, A., Pignatelli, M. & Bodmer, W. β-Catenin mutations in cell lines established from human colorectal cancers. Proceedings of the National Academy of Sciences 94, 10330–10334 (1997).

85 Iwao, K. et al. Activation of the β-catenin gene by interstitial deletions involving exon 3 in primary colorectal carcinomas without adenomatous polyposis coli mutations. Cancer research 58, 1021–1026 (1998).

86 Sparks, A. B., Morin, P. J., Vogelstein, B. & Kinzler, K. W. Mutational analysis of the APC/β-catenin/Tcf pathway in colorectal cancer. Cancer research 58, 1130–1134 (1998).

87 Huang, H. et al. APC mutations in sporadic medulloblastomas. The American journal of pathology 156, 433–437 (2000).

88 Zurawel, R. H., Chiappa, S. A., Allen, C. & Raffel, C. Sporadic medulloblastomas contain oncogenic β-catenin mutations. Cancer research 58, 896–899 (1998).

89 Dahmen, R. et al. Deletions of AXIN1, a component of the WNT/wingless pathway, in sporadic medulloblastomas. Cancer research 61, 7039–7043 (2001).

90 Fukuchi, T. et al. β-Catenin mutation in carcinoma of the uterine endometrium. Cancer research 58, 3526–3528 (1998).

91 Koch, A. et al. Childhood hepatoblastomas frequently carry a mutated degradation targeting box of the β-catenin gene. Cancer research 59, 269–273 (1999).

92 Wei, Y. et al. Activation of β-catenin in epithelial and mesenchymal hepatoblastomas. Oncogene 19, 498–504 (2000).

93 Schmidt, L. et al. Germline and somatic mutations in the tyrosine kinase domain of the MET proto-oncogene in papillary renal carcinomas. Nature genetics 16, 68–73 (1997).

94 Park, W. S. et al. Somatic mutations in the kinase domain of the Met/hepatocyte growth factor receptor gene in childhood hepatocellular carcinomas. Cancer research 59, 307–310 (1999).

95 Ma, P. C. et al. c-MET mutational analysis in small cell lung cancer: novel juxtamembrane domain mutations regulating cytoskeletal functions. Cancer research 63, 6272–6281 (2003).

96 McCoy, M. L., Mueller, C. R. & Roskelley, C. D. The role of the breast cancer susceptibility gene 1 (BRCA1) in sporadic epithelial ovarian cancer. Reproductive Biology and Endocrinology 1, 1–5 (2003).

97 Beer, S. et al. Developmental context determines latency of MYC-induced tumorigenesis. PLoS biology 2, e332 (2004).

98 Yang, Y. et al. The role of tripartite motif-containing 28 in cancer progression and its therapeutic potentials. Frontiers in Oncology 13, doi:10.3389/fonc.2023.1100134 (2023).

99 Wang, Y. et al. KAP1 is overexpressed in hepatocellular carcinoma and its clinical significance. International journal of clinical oncology 21, 927–933 (2016).

100 Varghese, S. et al. Site-specific gene expression profiles and novel molecular prognostic factors in patients with lower gastrointestinal adenocarcinoma diffusely metastatic to liver or peritoneum. Annals of surgical oncology 14, 3460–3471, doi:10.1245/s10434-007-9557-7 (2007).

101 Addison, J. B. et al. KAP1 promotes proliferation and metastatic progression of breast cancer cells. Cancer research 75, 344–355, doi:10.1158/0008-5472.Can-14-1561 (2015).

102 Czerwińska, P. et al. TRIM28 multi-domain protein regulates cancer stem cell population in breast tumor development. Oncotarget 8, 863 (2017).

103 Wei, C. et al. Tripartite motif containing 28 (TRIM28) promotes breast cancer metastasis by stabilizing TWIST1 protein. Sci Rep 6, 1–12 (2016).

104 Lin, L.-F. et al. Loss of ZBRK1 contributes to the increase of KAP1 and promotes KAP1-mediated metastasis and invasion in cervical cancer. PLoS One 8, e73033 (2013).

105 Yokoe, T. et al. KAP1 is associated with peritoneal carcinomatosis in gastric cancer. Annals of surgical oncology 17, 821–828 (2010).

106 Hu, M. et al. Expression of KAP1 in epithelial ovarian cancer and its correlation with drug-resistance. International journal of clinical and experimental medicine 8, 17308 (2015).

107 Cui, Y. et al. High levels of KAP1 expression are associated with aggressive clinical features in ovarian cancer. International journal of molecular sciences 16, 363–377, doi:10.3390/ijms16010363 (2014).

108 Qi, Z.-X. et al. TRIM28 as an independent prognostic marker plays critical roles in glioma progression. Journal of neuro-oncology 126, 19–26 (2016).

109 Jovčevska, I. et al. Differentially expressed proteins in glioblastoma multiforme identified with a nanobody-based anti-proteome approach and confirmed by OncoFinder as possible tumor-class predictive biomarker candidates. Oncotarget 8, 44141–44158, doi:10.18632/oncotarget.17390 (2017).

110 Chen, L. et al. Tripartite motif containing 28 (Trim28) can regulate cell proliferation by bridging HDAC1/E2F interactions. J Biol Chem 287, 40106–40118, doi:10.1074/jbc.M112.380865 (2012).

111 Bojkowska, K. et al. Liver-specific ablation of Krüppel-associated box–associated protein 1 in mice leads to male-predominant hepatosteatosis and development of liver adenoma. Hepatology 56, 1279–1290, 10.1002/hep.25767 (2012).

112 Cassano, M. et al. Polyphenic trait promotes liver cancer in a model of epigenetic instability in mice. Hepatology (Baltimore, Md.) 66, 235–251, doi:10.1002/hep.29182 (2017).

113 Song, T. et al. TRIM28 represses renal cell carcinoma cell proliferation by inhibiting TFE3/KDM6A-regulated autophagy. Journal of Biological Chemistry, 104621, 10.1016/j.jbc.2023.104621 (2023).

114 Park, H. H. et al. RIPK3 activation induces TRIM28 derepression in cancer cells and enhances the anti-tumor microenvironment. Mol Cancer 20, 107, doi:10.1186/s12943-021-01399-3 (2021).

115 Diets, I. J. et al. TRIM28 haploinsufficiency predisposes to Wilms tumor. International journal of cancer 145, 941–951, 10.1002/ijc.32167 (2019).

116 Sampath Kumar, A., et al. Loss of maternal Trim28 causes male-predominant early embryonic lethality. Genes & development 31, 12–17, doi:10.1101/gad.291195.116 (2017).

117 Messerschmidt, D. M. et al. Trim28 is required for epigenetic stability during mouse oocyte to embryo transition. Science (New York, N.Y.) 335, 1499–1502, doi:10.1126/science.1216154 (2012).

118 Lorthongpanich, C. et al. Single-cell DNA-methylation analysis reveals epigenetic chimerism in preimplantation embryos. Science (New York, N.Y.) 341, 1110–1112, doi:10.1126/science.1240617 (2013).

119 Blewitt, M. E. et al. An *N*-ethyl-*N*-nitrosourea screen for genes involved in variegation in the mouse. Proc. Natl. Acad. Sci. USA 102, 7629, doi:10.1073/pnas.0409375102 (2005).

120 Levene, H. in Contributions to Probability and Statistics 278–292 (1960).

121 Benjamini, Y. & Hochberg, Y. Controling the false discovery rate: a practical and powerful approach to multiple testing. J. Royal Stat. Soc. B 57, 289–300 (1995).

122 Scrucca, L., Fop, M., Murphy, T. B. & Raftery, A. E. mclust 5: Clustering, Classification and Density Estimation Using Gaussian Finite Mixture Models. The R journal 8, 289–317 (2016).

123 Lebret, R. et al. Rmixmod: The R Package of the Model-Based Unsupervised, Supervised, and Semi-Supervised Classification Mixmod Library. Journal of Statistical Software 67, 1–29, doi:10.18637/jss.v067.i06 (2015).

124 Hastie, T. & Tibshirani, R. in Monographs on Statistics & Applied Probability (Chapman and Hall/CRC, 1990).

125. Lenth, R., Singmann, H., Love, J., Buerkner, P. & Herve, M. (2019).

126 Searle, S. R., Speed, F. M. & Milliken, G. A. Population marginal means in the linear model: An alternative to least squares means. Am. Stat. 34, 216–221, doi:10.1080/00031305.1980.10483031 (1980).

127 Wilson, E. B. Probable inference, the law of succession, and statistical inference. J. Am. Stat. Assoc. 22, 209–212, doi:10.1080/01621459.1927.10502953 (1927).

128 Newcombe, R. G. Interval estimation for the difference between independent proportions: comparison of eleven methods. Stat Med 17, 873–890, doi:10.1002/(sici)1097-0258(19980430)17:8<873::aid-sim779>3.0.co;2-i (1998).

129 Newcombe, R. G. Two-sided confidence intervals for the single proportion: comparison of seven methods. Stat Med 17, 857–872, doi:10.1002/(sici)1097-0258(19980430)17:8<857::aid-sim777>3.0.co;2-e (1998).

130 Zhou, W. et al. DNA methylation dynamics and dysregulation delineated by high-throughput profiling in the mouse. Cell Genom. 2, doi:10.1016/j.xgen.2022.100144 (2022).

131 Zhou, W., Triche, T. J., Jr., Laird, P. W. & Shen, H. SeSAMe: reducing artifactual detection of DNA methylation by Infinium BeadChips in genomic deletions. Nucl. Acids Res. 46, e123, doi:10.1093/nar/gky691 (2018).

132 Apostle, S., Fagnocchi, L. & Pospisilik, J. A. SeSAMeStr: An automated pipeline for SeSAMe methylation array analysis (v1.0.0). Zenodo, 10.5281/zenodo.7510575 (2023).

133 Gu, Z., Eils, R. & Schlesner, M. Complex heatmaps reveal patterns and correlations in multidimensional genomic data. Bioinformatics 32, 2847–2849, doi:10.1093/bioinformatics/btw313 (2016).

134 Ritchie, M. E. et al. limma powers differential expression analyses for RNA-sequencing and microarray studies. Nucleic acids research 43, e47–e47, doi:10.1093/nar/gkv007 (2015).

135 Yu, G., Wang, L. G., Han, Y. & He, Q. Y. clusterProfiler: an R package for comparing biological themes among gene clusters. Omics 16, 284–287, doi:10.1089/omi.2011.0118 (2012).

136 Pletscher-Frankild, S., Pallejà, A., Tsafou, K., Binder, J. X. & Jensen, L. J. DISEASES: Text mining and data integration of disease–gene associations. Methods 74, 83–89, 10.1016/j.ymeth.2014.11.020 (2015).

137 Kuleshov, M. V. et al. Enrichr: a comprehensive gene set enrichment analysis web server 2016 update. Nucl. Acids. Res. 44, W90–97, doi:10.1093/nar/gkw377 (2016).

138 Cerami, E. et al. The cBio cancer genomics portal: an open platform for exploring multidimensional cancer genomics data. Cancer Discov. 2, 401–404, doi:10.1158/2159-8290.Cd-12-0095 (2012).

139 Gao, J. et al. Integrative analysis of complex cancer genomics and clinical profiles using the cBioPortal. Sci. Signal 6, pl1, doi:10.1126/scisignal.2004088 (2013).

