## Supplementary figures and tables for "Developmental priming of cancer susceptibility"

### Suppl.Fig.1

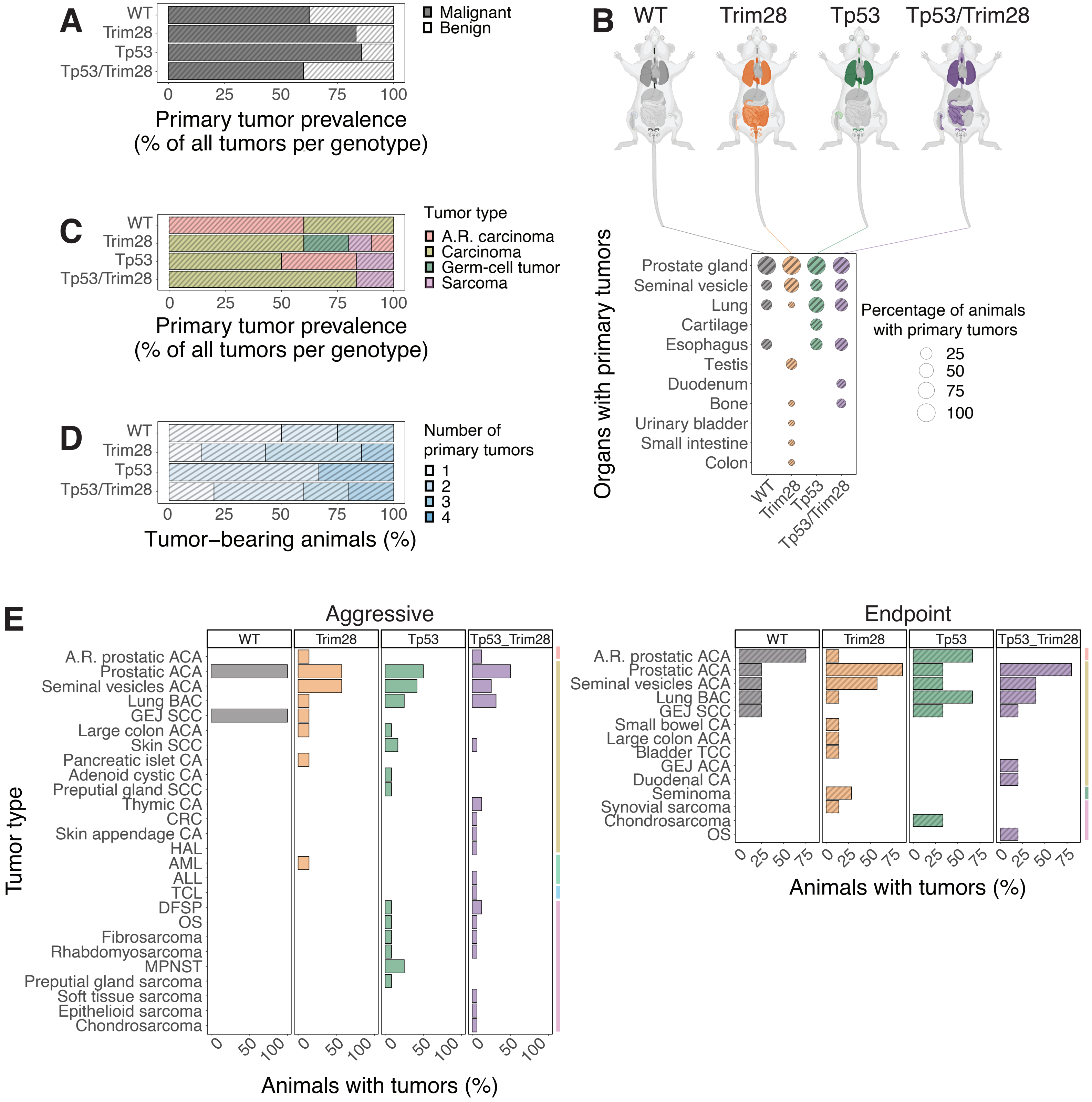

A
WT
Trim28
Tp53
Tp53/Trim28
0 25 50 75 100
Primary tumor prevalence
(% of all tumors per genotype)
Malignant
Benign
B
WT
Trim28
Tp53
Tp53/Trim28
Organs with primary tumors
Prostate gland
Seminal vesicle
Lung
Cartilage
Esophagus
Testis
Duodenum
Bone
Urinary bladder
Small intestine
Colon
Percentage of animals with primary tumors
25
50
75
100
C
Tumor type
A.R. carcinoma
Carcinoma
Germ-cell tumor
Sarcoma
D
Number of primary tumors
1
2
3
4
Tumor–bearing animals (%)
E
Aggressive
Endpoint
Tp53_Trim28
Tumor type
A.R. prostatic ACA
Prostatic ACA
Seminal vesicles ACA
Lung BAC
GEJ SCC
Large colon ACA
Skin SCC
Pancreatic islet CA
Adenoid cystic CA
Preputial gland SCC
Thymic CA
CRC
Skin appendage CA
HAL
AML
ALL
TCL
DFSP
OS
Fibrosarcoma
Rhabdomyosarcoma
MPNST
Preputial gland sarcoma
Soft tissue sarcoma
Epithelioid sarcoma
Chondrosarcoma
Small bowel CA
Bladder TCC
GEJ ACA
Duodenal CA
Seminoma
Synovial sarcoma
Animals with tumors (%)

**Supplementary Figure 1. *Trim28$^{+/D9}$* mice exhibit a novel multi-cancer syndrome. A)** Prevalence of *malignant* (black bars) or *benign* (white bars) *endpoint* tumors of for each genotype, expressed as percentage relative to the total number of tumors found in each genotype. N=37 (total tumors: 8 in WT, 12 in *Trim28$^{+/D9}$*, 7 in *Tp53$^{R270H/+}$*, 10 in *Tp53$^{R270H/+}$;Trim28$^{+/D9}$*). **B)** Tissues targeted by *malignant endpoint* tumors in the different genotypes. Top panel: mouse anatomy plots, with non-targeted in light-grey and targeted tissues colored according to the affected genotype: WT in black, *Trim28$^{+/D9}$* in orange, *Tp53$^{R270H/+}$* in green, *Tp53$^{R270H/+}$;Trim28$^{+/D9}$* in purple. Bottom panel: percentage of animals with specific organs targeted by *malignant endpoint* tumors in the different genotypes. N=19 (4 WT, 7 *Trim28$^{+/D9}$*, 3 *Tp53$^{R270H/+}$*, 5 *Tp53$^{R270H/+}$;Trim28$^{+/D9}$*). Created with BioRender.com. **C)** Prevalence of distinct *malignant endpoint* tumor types for each genotype, expressed as percentage relative to the total number of *malignant endpoint* tumors found in each genotype. N=27 (total *malignant endpoint* tumors: 5 in WT, 10 in *Trim28$^{+/D9}$*, 6 in *Tp53$^{R270H/+}$*, 6 in *Tp53$^{R270H/+}$;Trim28$^{+/D9}$*). **D)** Fraction of animals harboring 1 or multiple *malignant endpoint* tumors in the different genotypes. N=19 (4 WT, 7 *Trim28$^{+/D9}$*, 3 *Tp53$^{R270H/+}$*, 5 *Tp53$^{R270H/+}$;Trim28$^{+/D9}$*). **E)** Prevalence of distinct *malignant* tumor types for each genotype, divided into *aggressive* (left panel) and *endpoint* (right panel) and expressed as percentage relative to the total number of animals found with *aggressive* or *endpoint* tumors. N=60 (6 WT, 15 *Trim28$^{+/D9}$*, 17 *Tp53$^{R270H/+}$*, 22 *Tp53$^{R270H/+}$;Trim28$^{+/D9}$*). In both panels, colored bars on the right identify the main tumor type, as in Fig.1D (bottom panel) and S1C: age-related carcinoma in red; carcinoma in gold; leukemia in light green; lymphoma in light blue; sarcoma in pink; germ-cell tumors in dark green. N=60 (6 WT, 15 *Trim28$^{+/D9}$*, 17 *Tp53$^{R270H/+}$*, 22 *Tp53$^{R270H/+}$;Trim28$^{+/D9}$*).

### Suppl.Fig.2

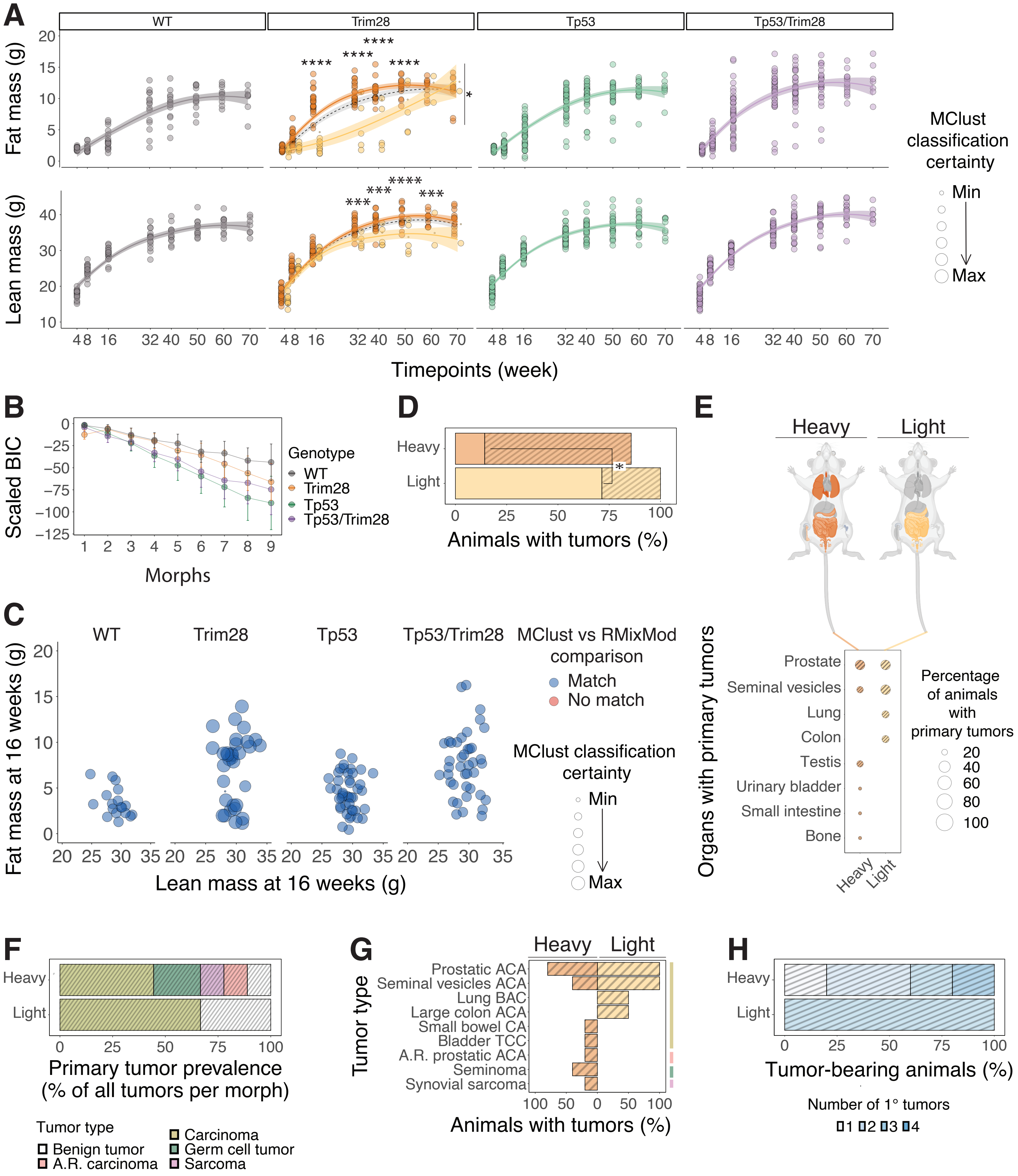
A
WT
Trim28
Tp53
Tp53/Trim28
Fat mass (g)
Lean mass (g)
Timepoints (week)
MClust classification certainty
Min
Max
B
Scaled BIC
Morphs
Genotype
WT
Trim28
Tp53
Tp53/Trim28
D
Heavy
Light
Animals with tumors (%)
E
Heavy
Light
Organs with primary tumors
Prostate
Seminal vesicles
Lung
Colon
Testis
Urinary bladder
Small intestine
Bone
Percentage of animals with primary tumors
C
Fat mass at 16 weeks (g)
Lean mass at 16 weeks (g)
MClust vs RMixMod comparison
Match
No match
MClust classification certainty
F
Primary tumor prevalence (% of all tumors per morph)
Tumor type
Benign tumor
A.R. carcinoma
Carcinoma
Germ cell tumor
Sarcoma
G
Tumor type
Prostatic ACA
Seminal vesicles ACA
Lung BAC
Large colon ACA
Small bowel CA
Bladder TCC
A.R. prostatic ACA
Seminoma
Synovial sarcoma
Animals with tumors (%)
H
Tumor-bearing animals (%)
Number of 1° tumors

**Supplementary Figure 2. TRIM28-dependent developmental heterogeneity primes cancer outcomes. A)** Scatter plots and smoothed conditional means (loess method) for *fat* (top panel) and *lean* (bottom panel) mass data for all the animals in the different genotypes at each analyzed timepoint (one dot represents one animal). The dot size indicates the classification certainty (probability) as calculated by MClust for each single animal. The different colors in the *Trim28*$^{+/D9}$ population represent the classification into 2 distinct clusters, while the other genotypes are better described by 1 cluster. Threshold of significance for padj<0.01 for GAM and emmeans analysis to compare overall slope differences and differences within each timepoint. *** p<0.001, *** p<0.0001. Top panel: p=0.0238 for *Trim28*$^{+/D9}$*-heavy* vs *-light* overall fat mass trajectories. Timepoint-specific comparisons: p<0.0001 at 16, 30, 40, and 50 weeks of age. Bottom panel: p= 0.0660 for *Trim28*$^{+/D9}$*-heavy* and *-light* overall lean mass trajectories. Timepoint-specific comparisons: p= 0.0002 at 32 weeks of age, p= 0.0001 at 40 weeks of age, p<0.0001 at 50 weeks of age, and p= 0.0009 at 60 weeks of age. N=137 (18 WT, 34 *Trim28*$^{+/D9}$, 44 *Tp53*$^{R270H/+}$, 41 *Tp53*$^{R270H/+}$*;Trim28*$^{+/D9}$). **B)** Mean and standard deviation of scaled Bayesian Information Criterion (BIC) determined for the 10 models tested by MClust for each cluster number option (morph) and genotype. The data are the same as in S2A. **C)** Fat and lean mass data at 16 weeks for all the animals in the different genotypes (each dot represents one animal). The dot size represents the classification certainty (probability) as calculated by MClust for each animal. A red dot would indicate an animal that is classified differently by Rmixmod and MClust. All the dots are blue, indicating 100% concordance of the 2 clustering methods. The data are the same as in Fig.2B. **D)** Proportion of *Trim28*$^{+/D9}$*-heavy* and *-light* animals affected by *malignant aggressive* (empty bars) or *endpoint* (patterned bars) tumors (expressed as percentage of animals developing tumors relative to the total animals in each population). Two-sample test for equality of proportions without continuity correction, significance for * p<0.05: *-heavy* vs *-light* with *aggressive malignant* tumors, p=0.0308, $\chi^2$=4.6667; *-heavy* vs *-light* with *malignant endpoint* tumors, p=0.1088, $\chi^2$=2.5714. N=14 (7 *-heavy*, 7 *-light*). **E)** Tissues targeted by *malignant endpoint* tumors in the different genotypes. Top panel: mouse anatomy plots, with non-targeted in light-grey and targeted tissues colored according to the affected morph: dark orange for *-heavy*, pale orange for *-light*. Bottom panel: percentage of animals with specific organs targeted by *malignant endpoint* tumors in the different morphs. N=7 (5 *-heavy*, 2 *-light*). **F)** Prevalence of distinct *endpoint* tumor types in *Trim28*$^{+/D9}$*-heavy* and *-light* animals (expressed as percentage relative to the total number of *endpoint* tumors found in each population). N=12 (total *endpoint* tumors, including *malignant* and *benign*, 7 in *-heavy* and 3 in *-light*). **G)** Distribution of *malignant endpoint* tumor types in *Trim28*$^{+/D9}$*-heavy* and *-light* animals, expressed as percentage of animals with a particular tumor type relative to the total number of animals with *malignant endpoint* tumors in each population. The colored bars on the right identify the main tumor type, as in S2F: carcinoma in gold; age-related carcinoma in red; germ cell tumor in dark green; sarcoma in pink. N=7 (5 *-heavy*, 2 *-light*). **H)** Fraction of *Trim28*$^{+/D9}$*-heavy* and *-light* animals that died at the endpoint of the study harboring 1 or multiple *malignant endpoint* tumors. N=7 (5 *-heavy*, 2 *-light*).

### Suppl.Fig.3

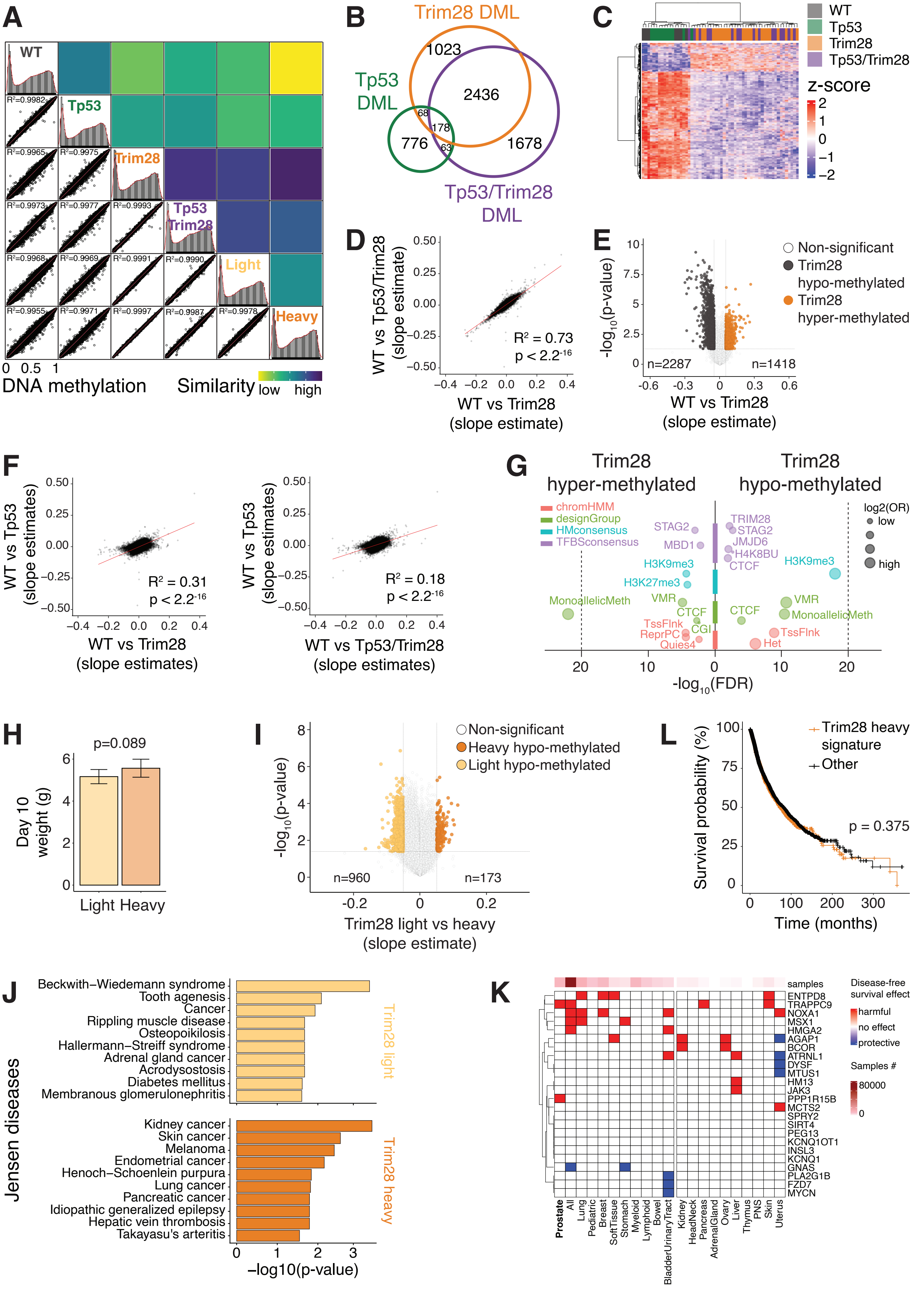
A
WT
Tp53
Trim28
Tp53 Trim28
Light
Heavy
R²=0.9982
R²=0.9965
R²=0.9975
R²=0.9973
R²=0.9977
R²=0.9993
R²=0.9968
R²=0.9969
R²=0.9991
R²=0.9990
R²=0.9955
R²=0.9971
R²=0.9997
R²=0.9987
R²=0.9978
0 0.5 1
DNA methylation
Similarity low high
B
Trim28 DML
Tp53 DML
Tp53/Trim28 DML
1023
2436
68
178
776
63
1678
C
WT
Tp53
Trim28
Tp53/Trim28
z-score
2
1
0
−1
−2
D
WT vs Tp53/Trim28 (slope estimate)
WT vs Trim28 (slope estimate)
R² = 0.73
p < 2.2⁻¹⁶
E
-log10(p-value)
WT vs Trim28 (slope estimate)
n=2287
n=1418
Non-significant
Trim28 hypo-methylated
Trim28 hyper-methylated
F
WT vs Tp53 (slope estimates)
WT vs Trim28 (slope estimates)
R² = 0.31
p < 2.2⁻¹⁶
WT vs Tp53/Trim28 (slope estimates)
R² = 0.18
p < 2.2⁻¹⁶
G
Trim28 hyper-methylated
Trim28 hypo-methylated
chromHMM
designGroup
HMconsensus
TFBSconsensus
log2(OR)
low
high
STAG2
MBD1
H3K9me3
H3K27me3
MonoallelicMeth
VMR
CTCF
CGI
TssFlnk
ReprPC
Quies4
TRIM28
STAG2
JMJD6
H4K8BU
CTCF
H3K9me3
VMR
MonoallelicMeth
CTCF
TssFlnk
Het
-log10(FDR)
H
p=0.089
Day 10 weight (g)
Light Heavy
I
-log10(p-value)
Non-significant
Heavy hypo-methylated
Light hypo-methylated
n=960
n=173
Trim28 light vs heavy (slope estimate)
L
Trim28 heavy signature
Other
Survival probability (%)
p = 0.375
Time (months)
J
Jensen diseases
Beckwith–Wiedemann syndrome
Tooth agenesis
Cancer
Rippling muscle disease
Osteopoikilosis
Hallermann–Streiff syndrome
Adrenal gland cancer
Acrodysostosis
Diabetes mellitus
Membranous glomerulonephritis
Trim28 light
Kidney cancer
Skin cancer
Melanoma
Endometrial cancer
Henoch–Schoenlein purpura
Lung cancer
Pancreatic cancer
Idiopathic generalized epilepsy
Hepatic vein thrombosis
Takayasu's arteritis
Trim28 heavy
−log10(p-value)
K
samples
ENTPD8
TRAPPC9
NOXA1
MSX1
HMGA2
AGAP1
BCOR
ATRNL1
DYSF
MTUS1
HM13
JAK3
PPP1R15B
MCTS2
SPRY2
SIRT4
PEG13
KCNQ1OT1
INSL3
KCNQ1
GNAS
PLA2G1B
FZD7
MYCN
Prostate
All
Lung
Pediatric
Breast
SoftTissue
Stomach
Myeloid
Lymphoid
Bowel
BladderUrinaryTract
Kidney
HeadNeck
Pancreas
AdrenalGland
Ovary
Liver
Thymus
PNS
Skin
Uterus
Disease-free survival effect
harmful
no effect
protective
Samples #
80000
0

**Supplementary Figure 3. *Trim28$^{+/D9}$*-dependent cancer susceptibility states are distinguished by distinct early-life epigenomes that are enriched for epigenetic regulators and bona fide oncogenes. A)** On the right, correlation plots comparing the global DNA methylation on all available probes among all genotypes, and the *Trim28$^{+/D9}$-heavy* and *-light* morphs as sum of squared residuals from linear regressions. On the left, density plots of beta values distribution for all genotypes/trajectories. $R^2$ values derive from Pearson correlations. N=58 (7 WT, 24 *Trim28$^{+/D9}$*, 11 *Tp53$^{R270H/+}$*, 16 *Tp53$^{R270H/+}$;Trim28$^{+/D9}$*). **B)** Overlap of DMLs, between *Trim28$^{+/D9}$*, *Tp53$^{R270H/+}$;Trim28$^{+/D9}$*, *Tp53$^{R270H/+}$* and WT animals. N=58 (7 WT, 24 *Trim28$^{+/D9}$*, 11 *Tp53$^{R270H/+}$*, 16 *Tp53$^{R270H/+}$;Trim28$^{+/D9}$*). **C)** Heatmap of the z-score of log1p transformed beta values of the differentially methylated probes in all genotypes relative to WT animals. Effect size cut-off = 0.1; Benjamini–Hochberg adjusted p-value cut-off=0.05. N=58 (7 WT, 24 *Trim28$^{+/D9}$*, 11 *Tp53$^{R270H/+}$*, 16 *Tp53$^{R270H/+}$;Trim28$^{+/D9}$*). **D)** Correlation plot comparing the DNA methylation levels (slope estimate) on all available probes in WT vs *Tp53$^{R270H/+}$;Trim28$^{+/D9}$* and WT vs *Trim28$^{+/D9}$* animals. $R^2$ value=0.73; $p<2.2^{-16}$ (Pearson correlation). N=58 (7 WT, 24 *Trim28$^{+/D9}$*, 11 *Tp53$^{R270H/+}$*, 16 *Tp53$^{R270H/+}$;Trim28$^{+/D9}$*). **E)** Volcano plot of the differential DNA methylation levels (slope estimate) in WT vs *Trim28$^{+/D9}$* animals. N=31 (7 WT vs 24 *Trim28$^{+/D9}$*). Black and orange dots represent hypo- and hyper-methylated probes in *Trim28$^{+/D9}$* animals, respectively. White dots are non-significant DMLs (estimate and p-value cut-off=0.05). **F)** Correlation plots comparing the DNA methylation levels (slope estimate) on all available probes in WT vs *Tp53$^{R270H/+}$* (left panel) and WT vs *Tp53$^{R270H/+}$;Trim28$^{+/D9}$* animals (right panel). $R^2$ values=0.31 and =0.18; $p<2.2^{-16}$ (Pearson correlation). N=58 (7 WT, 24 *Trim28$^{+/D9}$*, 11 *Tp53$^{R270H/+}$*, 16 *Tp53$^{R270H/+}$;Trim28$^{+/D9}$*). **G)** Enrichment plots of features for *Trim28$^{+/D9}$*-dependent differentially methylated probes. False Discovery Rate (FDR) cut-off=0.01 (one-tailed Fisher's exact test). **H)** Weight at day 10 of the *Trim28$^{+/D9}$-light* vs *-heavy* animals used for the DNA methylation array experiments. N=24 (15 *Trim28$^{+/D9}$-heavy* vs 9 *Trim28$^{+/D9}$-light*). P-value=0.089 (unpaired two-samples Wilcoxon test). **I)** Volcano plot of the differential DNA methylation levels (slope estimate) in *Trim28$^{+/D9}$-heavy* vs *-light* animals. Dark and pale orange dots represent hyper- and hypo-methylated probes in *-light* animals, respectively (estimate and p-value cut-off=0.05). White dots are non-significant DMLs. N=24 (15 *Trim28$^{+/D9}$-heavy* vs 9 *Trim28$^{+/D9}$-light*). **J)** Enrichments of the genes enriched by hypo-methylated probes in *Trim28$^{+/D9}$-light* (pale orange) vs *-heavy* animals (dark orange) on the Jensen DISEASES database (https://diseases.jensenlab.org/Search). P-value cut-off = 0.05. **K)** Heatmap of the effects on disease-free survival probability of mutations in the specified genes and tumor tissues. The analysis includes all samples from TCGA and non-TCGA studies with no overlapping samples, from cBioPortal (N=69223 samples). Tumor tissues are separated in two main branches according to samples numerosity (left: >3000 samples; right: <3000 samples) and ordered as in Fig.3F. **L)** The Kaplan-Meier survival probability as percentage of the total population and the time of survival in month for all TCGA PanCancer Atlas patients mutated in genes from the *Trim28$^{+/D9}$-heavy* hypo-methylated signature (dark orange, n=1865) compared to individuals not mutated in the same genes (black, n=9085). The two do not exhibit differential survival (log-rank test, p=0.375). Total cases analyzed = 10967.

### Appendix 1

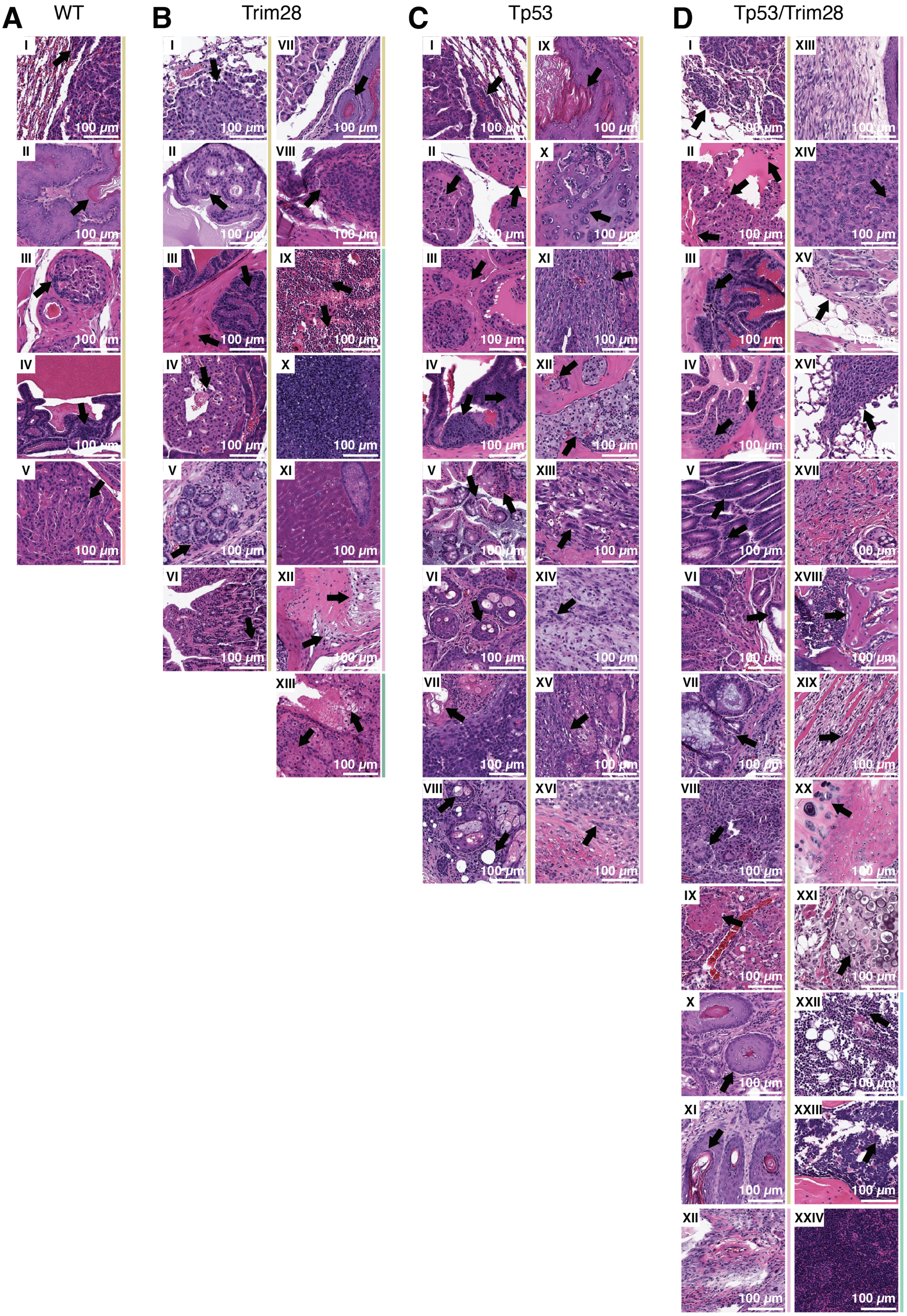

**Appendix 1. Representative images of histological examples of tumor types and targeted tissue for all genotypes and timepoints.** In all panels, arrows indicate main features of each tumor type. Colored bars on the right identify the main tumor type, as in Fig.1F, bottom panel: age-related carcinoma in red; carcinoma in gold; leukemia in light green; lymphoma in light blue; sarcoma in pink; germ-cell tumors in dark green. **A)** Representative images for WT animals. I- Bronchoalveolar carcinoma (BAC). II- Gastro-esophageal junction squamous cell carcinoma (SCC). III- Prostatic carcinoma (PCa). IV- Seminal vesicles carcinoma. V- Age-related PCa. **B)** Representative images for *Trim28*$^{+/D9}$ animals. I- BAC. II- PCa. III- Seminal vesicles carcinoma. IV- Pancreatic carcinoma. V- Large colon ACA. VI- Small bowel carcinoma. VII- Gastro-esophageal junction SCC. VIII- Bladder transitional cell carcinoma (TCC). IX- Acute myeloid leukemia (AML) affecting the bone marrow. X- AML spreading to the spleen. XI- AML spreading to the liver. XII- Synovial sarcoma. XIII- Seminoma. **C)** Representative images for *Tp53*$^{R270H/+}$ animals. I- Bronchoalveolar carcinoma (BAC). II- Age-related PCa. III- PCa. IV- Seminal vesicles carcinoma. V- Large colon adenocarcinoma (ACA). VI- Skin SCC. VII- Preputial glands SCC. VIII- Adenoid cystic carcinoma. IX- Gastro-esophageal junction SCC. X- Osteosarcoma (OS). XI- Fibrosarcoma. XII- Chondrosarcoma. XIII- Rhabdomyosarcoma. XIV- Malignant Peripheral Nerve Sheath Tumor (MPNST). XV- Sarcoma affecting the preputial glands. XVI- Dermatofibrosarcoma protuberans (DFSP). **D)** Representative images for *Tp53*$^{R270H/+}$*;Trim28*$^{+/D9}$ animals. I- BAC. II- PCa. III- Seminal vesicles carcinoma. IV- Age-related PCa. V- Gastro-esophageal junction ACA. VI- Duodenal carcinoma. VII- Colorectal carcinoma (CRC). VIII- Thymic carcinoma. IX- Hepatoid adenocarcinoma of the lung (HAL). X- Gastro-esophageal junction SCC. XI- Skin SCC. XII- DFSP. XIII- Soft tissue sarcoma. XIV- Epithelioid fibrohistiocytic sarcoma. XV- Skin appendage tumor. XVI- Metastatic fibrosarcoma to the lungs. XVII- Fibrosarcoma. XVIII- OS. XIX- Rhabdomyosarcoma. XX- Chondrosarcoma in the bone. XXI- Chondrosarcoma spreading to the soft tissue. XXII- T-cell lymphoma (TCL). XXIII- Acute lymphoblastic leukemia (ALL) affecting the bone marrow. XXIV- ALL spreading to the spleen.

#### SUPPLEMENTARY TABLES

##### Table 1 – Genotyping primers

| Target | Forward primer | Reverse primer |
|---|---|---|
| *Trim28* | CATGGCCATTGTCAAGGTAAG | AGGAGAACACGCTCACATTTC |
| *Tp53* | ATGCGACTCTCCAGCCTTGGTA | TTGGGCTTAGGGACGTCTCTTATC |

##### Table 2 – Genotyping PCR conditions

| Target | Initial denaturation | Denaturation (x40 cycles) | Annealing (x40 cycles) | Extension (x40 cycles) | Final extension | Hold |
|---|---|---|---|---|---|---|
| *Trim28* | 95 °C for 3 minutes | 95 °C for 30 seconds | 59 °C for 30 seconds | 72 °C for 1 minute | 72 °C for 5 minutes | 4 °C infinite |
| *Tp53* | 95 °C for 3 minutes | 95 °C for 30 seconds | 63 °C for 30 seconds | 72 °C for 1 minute | 72 °C for 5 minutes | 4 °C infinite |

##### Table 3 – Genotyping restriction conditions

| Target | Digestion | Enzyme inactivation | Hold |
|---|---|---|---|
| *Trim28* | 37 °C for 30 minutes (XceI/NspI) | 65 °C for 10 minutes | 4 °C infinite |
| *Tp53* | 37 °C for 30 minutes (MslI) | 80°C for 20 minutes | 4 °C infinite |
